# TRPV4-Expressing Tissue-Resident Macrophages Regulate the Function of Collecting Lymphatic Vessels via Thromboxane A2 Receptors in Lymphatic Muscle Cells

**DOI:** 10.1101/2024.05.21.595189

**Authors:** Mary E. Schulz, Victoria L. Akerstrom, Kejing Song, Sarah E. Broyhill, Min Li, Michelle D. Lambert, Tatia B. Goldberg, Raghu P. Kataru, Jinyeon Shin, Stephen E. Braun, Charles E. Norton, Rafael S. Czepielewski, Babak J. Mehrara, Timothy L. Domeier, Scott D. Zawieja, Jorge A. Castorena-Gonzalez

## Abstract

**Rationale:** TRPV4 channels are critical regulators of blood vascular function and have been shown to be dysregulated in many disease conditions in association with inflammation and tissue fibrosis. These are key features in the pathophysiology of lymphatic system diseases, including lymphedema and lipedema; however, the role of TRPV4 channels in the lymphatic system remains largely unexplored. TRPV4 channels are calcium permeable, non-selective cation channels that are activated by diverse stimuli, including shear stress, stretch, temperature, and cell metabolites, which may regulate lymphatic contractile function.

**Objective:** To characterize the expression of TRPV4 channels in collecting lymphatic vessels and to determine the extent to which these channels regulate the contractile function of lymphatics.

**Methods and Results:** Pressure myography on intact, isolated, and cannulated lymphatic vessels showed that pharmacological activation of TRPV4 channels with GSK1016790A (GSK101) led to contractile dysregulation. The response to GSK101 was multiphasic and included, 1) initial robust constriction that was sustained for ≥1 minute and in some instances remained for ≥4 minutes; and 2) subsequent vasodilation and partial or complete inhibition of lymphatic contractions associated with release of nitric oxide. The functional response to activation of TRPV4 channels displayed differences across lymphatics from four anatomical regions, but these differences were consistent across different species (mouse, rat, and non-human primate). Importantly, similar responses were observed following activation of TRPV4 channels in arterioles. The initial and sustained constriction was prevented with the COX inhibitor, indomethacin. We generated a *controlled* and *spatially defined* single-cell RNA sequencing (scRNAseq) dataset from intact and microdissected collecting lymphatic vessels. Our data uncovered a subset of macrophages displaying the highest expression of *Trpv4* compared to other cell types within and surrounding the lymphatic vessel wall. These macrophages displayed a transcriptomic profile consistent with that of tissue-resident macrophages (TRMs), including differential expression of *Lyve1*, *Cd163*, *Folr2*, *Mrc1*, *Ccl8*, *Apoe*, *Cd209f*, *Cd209d*, and *Cd209g*; and at least half of these macrophages also expressed *Timd4.* This subset of macrophages also highly expressed *Txa2s*, which encodes the thromboxane A2 (TXA2) synthase. Inhibition of TXA2 receptors (TXA2Rs) prevented TRPV4-mediated contractile dysregulation. TXA2R activation on LMCs caused an increase in mobilization of calcium from intracellular stores through Ip3 receptors which promoted store operated calcium entry and vasoconstriction.

**Conclusions:** Clinical studies have linked cancer-related lymphedema with an increased infiltration of macrophages. While these macrophages have known anti-inflammatory and pro-lymphangiogenic roles, as well as promote tissue repair, our results point to detrimental effects to the pumping capacity of collecting lymphatic vessels mediated by activation of TRPV4 channels in macrophages. Pharmacological targeting of TRPV4 channels in LYVE1-expressing macrophages or pharmacological targeting of TXA2Rs may offer novel therapeutic strategies to improve lymphatic pumping function and lymph transport in lymphedema.

**Graphical Abstract:** 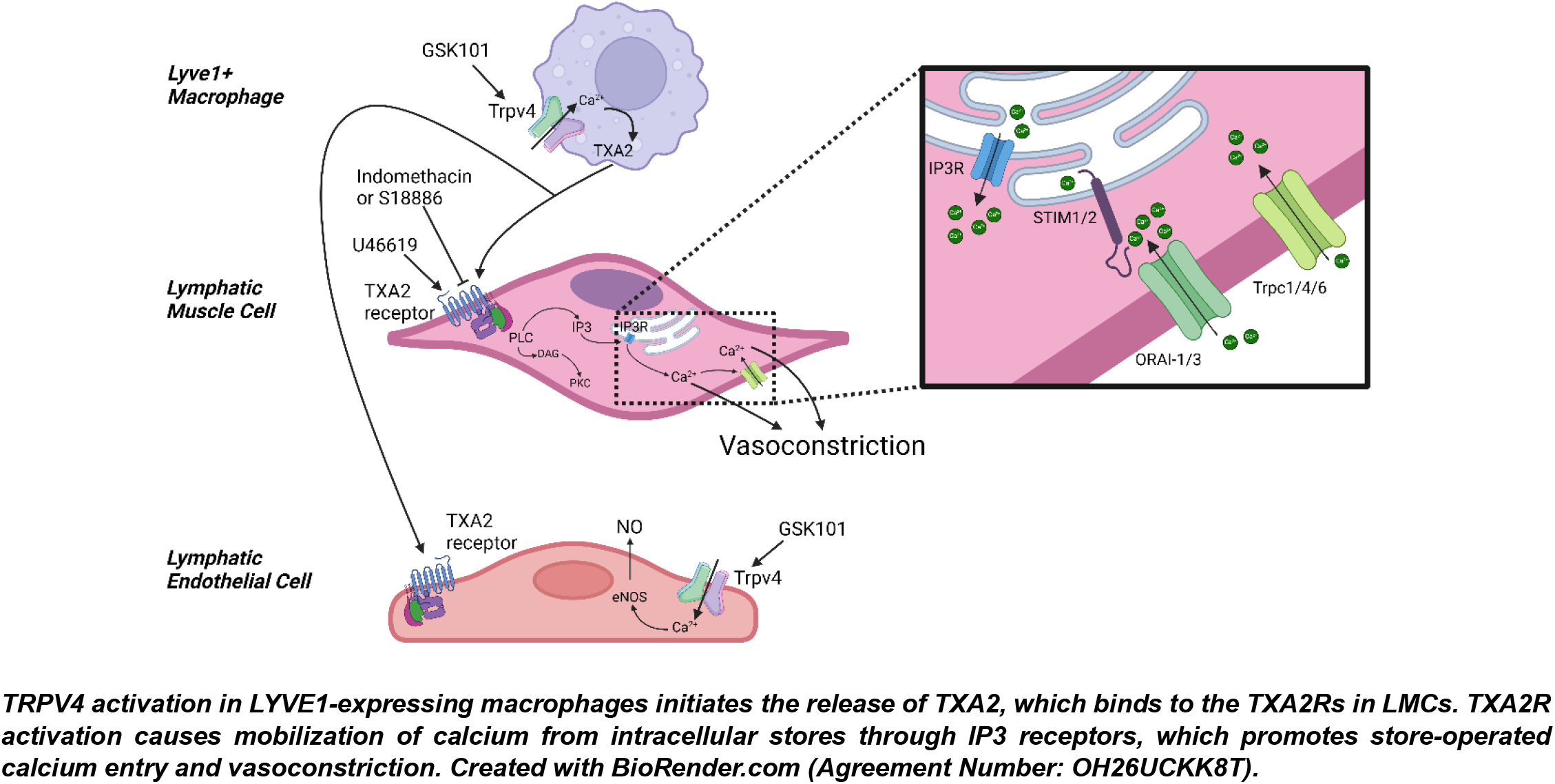

## Introduction

The lymphatic system plays fundamental roles in maintaining fluid homeostasis and immunity. Critical to these roles is the efficient transport of lymph (a fluid rich in proteins, lipids, immune cells, pathogens, cellular waste products, etc.) from the interstitial spaces back into the central veins. Following uptake by the lymphatic capillaries, lymph is transported through collecting lymphatic vessels by intrinsic forces (i.e., spontaneous, propulsive contractions), extrinsic forces (i.e., compression of lymphatic networks by external mediators including skeletal muscle contraction), and unidirectional intraluminal valves that ensure net forward flow. Dysfunction of the lymphatic system results in abnormal accumulation of fluid in the interstitial spaces throughout the body known as lymphedema. Some of the major consequences for patients afflicted with lymphedema and other lymphatic system diseases include pain, immobility, increased risk for infections, fibrosis, tissue damage, adipose tissue deposition, and impaired immune responses^1–4^. At present, there are no established pharmacological treatments or cures for lymphedema, only a single drug is currently being investigated for treatment of upper arm lymphedema^5,6^.

Relevant to the central topic of this study, signaling through Transient Receptor Potential Vanilloid Type 4 (TRPV4) channels has been established as a major regulator of vascular function^7–13^. TRPV4 channels are calcium-permeable, non-selective cation channels that are susceptible to activation by a variety of stimuli, including mechanical stimuli (e.g., shear stress and stretch), cell metabolites, chemical/pharmacological agents, temperature, changes in pH, hypoxia, as well as cell swelling and osmotic stress among others^9,14,15^.

Dysregulated signaling through TRPV4 channels has been linked to a variety of inflammatory diseases, including lung ischemia-reperfusion injury and other lung diseases^16–18^, cerebral edema following traumatic brain injury^19^, ocular vascular diseases^20–22^, obesity^23,24^, Parkinson’s disease^25^, various forms of cardiac dysfunction^26–30^, etc. Importantly, in all these scenarios, aberrant signaling through TRPV4 channels is associated with inflammation and tissue fibrosis, which are key features in the pathophysiology of lymphatic system disease, including lymphedema and lipedema^1,31^. The unexplored possibility is that dysregulated signaling through TRPV4 channels may be present in patients afflicted with lymphatic system diseases in association with chronic inflammation, fibrosis, and tissue damage/injury. Understanding how TRPV4 channels regulate the function of lymphatic vessels may offer key insights into novel therapeutic strategies. At present, the expression of TRPV4 channels within and surrounding the wall of collecting lymphatic vessels, and the extent to which activation of TRPV4 channels can regulate the different aspects of lymphatic vessel function (i.e., contractility, valve competency, and permeability) remains largely unexplored. Previous work demonstrated that in contrast to resistance arteries, where activation of endothelial TRPV4 channels promotes vasodilation mediated by endothelium-derived hyperpolarization^32,33^, stimulation of TRPV4 channels in mouse lymphatic endothelial cells (LECs) results in influx of calcium and depolarization of the cell membrane^34^. Utilizing rat diaphragmatic lymphatic vessels, Solari et al. demonstrated important roles for TRPV4 channels in modulating lymphatic contractility by means of sensing changes in temperature and osmolarity^35,36^. Recently, TRPV4 channels were shown to play a critical role in flow-mediated downregulation of the contractile frequency of rat mesenteric lymphatic vessels in association with production of nitric oxide and vasodilatory prostanoids^37^.

In this study, we present one of the first single-cell RNA sequencing (scRNAseq) datasets generated from isolated lymphatic vessels^38^. Our study presents a *controlled* and *spatially defined* scRNAseq dataset that included collecting lymphatic vessels from multiple anatomical regions and compared both sexes. In addition, we employed pressure myography experiments on intact, cannulated, and pressurized collecting lymphatic vessels to determine the specific cell types expressing TRPV4 channels and to identify and validate the different mechanisms by which these channels can regulate the contractile function of collecting lymphatic vessels. We found that in addition to LECs, TRPV4 channels are highly expressed in a subset of perivascular macrophages. Our scRNAseq data revealed that these are LYVE1-expressing macrophages and display a tissue-resident macrophage (TRM) profile^39–42^ that included, but are not limited to, differential expressions of *Cd163*, *Folr2*, *Mrc1*, *Ccl8*, *Apoe*, *Cd209f*, *Cd209d*, and *Cd209g*. At least half of the macrophages in this subpopulation also expressed another maker of TRMs, *Timd4*. Importantly, in a clinical study, biopsies from patients with upper-extremity breast cancer-related lymphedema showed a significantly increased infiltration of macrophages compared to controls; furthermore, the characteristics of these macrophages were consistent with M2-polarization^43^. It is known that these macrophages have anti-inflammatory and angiogenic functions and are involved in tissue remodeling and repair^44,45^. We demonstrate that signaling through TRPV4 channels leads to the activation of macrophages and subsequent secretion of Thromboxane A2 (TXA2), leading to major impairment of lymphatic contractility, which was significantly more profound (i.e., greater functional impairment) in lymphatics from male mice. Similar responses were also observed in arterioles. Our findings suggest that in lymphedema patients, lymphatic contractile dysfunction and impaired lymph transport could result from the increased infiltration of macrophages and the overall increased activity of TRPV4 channels in association with breast cancer-related therapies, injury/trauma, and/or chronic inflammation. Targeting TRPV4 channels in LYVE1-expressing macrophages and/or TXA2 receptors (TXA2Rs) in lymphatic muscle cells (LMCs) may offer novel therapeutic strategies to improve lymphatic contractility and lymph transport in lymphedema.

## Methods

### Animals, Microdissection of Collecting Lymphatic Vessels, and Pressure Myography

C57BL/6J (Cat. No.: 000664, WT) male and female mice were purchased from The Jackson Laboratory (Bar Harbor, ME) at 10-12 weeks of age. Mice were housed in a temperature-controlled room under a 12-hour light/dark cycle and had free access to regular food and water. Mice were allowed to acclimate for at least 2 weeks prior to experimentation. All mice were used between 3-6 months of age. Mice were weighed, anesthetized using isoflurane, and prepared for tissue dissection (see details for dissection of lymphatics from specific anatomical regions below). Sprague Dawley rats were provided by the *Tulane Department of Comparative Medicine*. These rats were purchased from Charles River at 6 weeks of age and used for experiments between 4-6 months of age. All mouse and rat experiments were performed under approved IACUC protocols number 1884 and 1602 respectively.

#### Dissection of popliteal lymphatic vessels

Starting with a mouse or rat in the prone position, a distal cut was initiated near the ankle and extended proximally to carefully expose the saphenous vein. The popliteal afferent lymphatic vessels run parallel to the saphenous vein, with one lymphatic vessel on each side of the vein. Krebs buffer was added to the popliteal region, and the popliteal lymphatic vessels were carefully dissected from the mouse, along with excess connective tissue and adipose tissue. Each tissue was then pinned down in a dissection chamber containing Krebs buffer to isolate the popliteal lymphatic away from connective tissue and adipose. Once isolated, vessels that were prepared for pressure myography experiments would be cannulated on glass micropipettes and further cleaned of adipose if necessary^46,47^.

#### Dissection of inguinal axillary lymphatic vessels (IALVs)

Starting with a mouse or rat in the prone position, dissection was initiated near the base of the tail and extended cephalad to the scapula. A small cut was made laterally to flay open the area. This tissue was pinned down on a dissection board, and Krebs buffer was added to keep the area moist. The IALVs run between two fat pads (one in the inguinal region and one in the axilla) and interconnect both the inguinal and the axillary lymph nodes. The lymphatic vessels, along with the two fat pads, were excised from the mouse and then pinned down in a dissection chamber with Krebs buffer to isolate the lymphatic vessel from the connective tissue and surrounding adipose prior to cannulation for pressure myography experiments^47^.

#### Dissection of mesenteric lymphatic vessels

Starting with a mouse or rat in the supine position, the entire mesentery was dissected from the rodent and placed in a dissection dish with Krebs buffer. Once the mesenteric lymph nodes were located, lymphatic vessels stemming from the lymph nodes were identified and carefully dissected. Mesenteric lymphatic vessels were then transferred into a dissection chamber, pinned down, and cleaned of residual adipose tissue^47,48^. Dissection of mesenteric tissue from rhesus macaques was done by the Tulane National Primate Research Center necropsy team.

#### Dissection of superficial cervical lymphatic vessels

Starting with a mouse or rat in the supine position, dissection was started centrally, near the jugular notch, extended up to the chin, and advanced laterally along the cheek, adding Krebs buffer as needed. Another lateral cut was made from the jugular notch along the clavicle to expose the superficial cervical lymph nodes. These lymph nodes were excised along with the blood vessels and lymphatic vessels by cutting from the lymph nodes up towards the masseter muscle and cheek. After excision, the lymph nodes and connected vessels were pinned down in a dissection dish with Krebs buffer to further clean and isolate the superficial cervical lymphatic vessels prior to cannulation^47^.

In preparation for pressure myography experiments, lymphatic vessels were carefully dissected and placed in Krebs buffer. Vessels were then transferred into an observation chamber containing the same buffer and cannulated on glass micropipettes, pressurized at 3 cmH_2_O, and equilibrated with perfusion at 37°C for 30 minutes before experimentation. The contractile activity of these lymphatic vessels was assessed under no-flow conditions. OB1 MK3 microfluidic flow control systems (Elveflow, Paris) were utilized for fine control of intraluminal pressures. Brightfield videos of contractions were recorded at 30 fps, and space-time maps (STMs) were generated as described in the following section.

### Generation of Space Time Maps (STMs) and Spectrograms

Brightfield videos of cannulated lymphatic vessels were recorded and processed using custom-made Python programs^46,49^. STMs were generated by tracking outside diameter as a function of time at every position along the entire lymphatic segment using previously developed methodologies. In these maps, diameter values are encoded in grayscale, where darker grays indicate smaller diameter (e.g., during each spontaneous contraction), and lighter gray shades indicate larger diameter (e.g., during diastole). For image representation, values are then normalized to span the entire range of an 8-bit scale using the largest and smallest diameter recorded values. Each darker band extending along the entire vertical axis indicates a contraction wave that propagated through the whole lymphatic segment. For further analysis, we generated two-dimensional frequency maps as a function of time, also known as spectrograms.

### Single Cell RNA Sequencing and Data Analysis

#### Sample Preparation

The single cell RNA sequencing dataset was generated from C57BL/6J mice (n=4 for male and female respectively). These mice were purchased from The Jackson Laboratory (Bar Harbor, ME) at 10-12 weeks of age and utilized for experiments after a 2-week acclimation period. Lymphatic vessels were carefully dissected as described in the previous section and placed in ice-cold low-calcium physiological saline solution (PSS) containing 1 mg/mL trypsin inhibitor, 0.1 mg/mL DNase I, 1 mM DTT, and 0.02 U/µL RNase inhibitor. For each mouse, lymphatic vessels from four anatomical regions (i.e., popliteal afferent, inguinal axillary, mesenteric, and superficial cervical) were combined into a single sample. A two-step tissue dissociation process was utilized. Tissues were transferred into enzymatic solution part 1 containing 1 mg/mL trypsin inhibitor, 0.1 mg/mL DNase I, 1 mM DTT, 0.02 U/µL RNase inhibitor, and 1 mg/mL papain in low-calcium PSS, and incubated at 37°C for 30 min with intermittent shaking every 9 minutes for 1 minute at 800 rpm. Meanwhile, enzyme solution part 2 was prepared: 1 mg/mL trypsin inhibitor, 0.1 mg/mL DNase I, 1 mM DTT, 0.02 U/µL RNase inhibitor, 1 mg/mL papain, 2 mg/mL collagenase A, and 3 mg/mL dispase II in low-calcium PSS. After the 30-minute incubation in enzyme solution part 1, vessels were transferred into enzyme solution part 2 and incubated for an additional 30 min at 37°C with intermittent shaking every 7 minutes for 1 minute at 800 rpm. Single-cell suspensions were ensured via pipetting up and down with a P200 pipette and a fine-tip Pasteur glass pipette. The volume of the single cell suspensions was doubled with cold low-calcium PSS to lower concentration of enzymes and stop/minimize enzymatic activity, and then cell suspensions were centrifuged at 600xg for 8 min at 4°C. Following centrifugation, supernatants were carefully removed, and cell pellets were gently resuspended with 200 µL sterile PBS containing 0.04% ultra-pure BSA. Cells were strained into a 1.5 mL LoBind centrifuge tube and cell counts and viability were assessed.

#### Feature Barcode Technology and Single Cell RNA Sequencing

Cell numbers and viabilities were validated by Cellometer Automated Cell Counter (Nexcelom Bioscience, Lawrence, MA, USA) prior to scRNAseq library preparation. For feature barcode technology and 10x scRNAseq assay, individual live cells per sample were targeted using feature barcode oligonucleotide conjugated to a lipid. Labeled cells from each sample were pooled and 10x Single Cell 3’ RNAseq technology was applied to generate both cell multiplexing libraries and cDNA libraries. Briefly, single cells from each sample were labeled with unique cell multiplexing oligo (CMO) provided by 10x Genomics (10X Genomics Inc, CA). 5,000 labeled cells from each sample were pooled and partitioned into nanoliter-scale Gel Beads-In-EMulsion (GEMs). Feature barcode molecules were captured directly by oligonucleotides presented on the Gel Beads inside a GEM during GEM-RT. Cell multiplexing libraries comprising dual index NN set A (10X Genomics Inc, CA) were generated by index PCR amplification. Meanwhile, full-length barcoded cDNAs were also generated and amplified by PCR to obtain sufficient mass for library construction. Following enzymatic fragmentation, end-repair, A-tailing, and adaptor ligation, single cell 3’ libraries comprising dual index TT set A (10X Genomics Inc, CA) were generated. Library quality controls were performed by using Agilent High Sensitive DNA kit with Agilent 2100 Bioanalyzer (Agilent Technologies, Palo Alto, CA, USA) and quantified by Qubit 2.0 fluorometer (Thermo Fisher Scientific). Pooled libraries at a final concentration of 750pM were sequenced with paired end dual index configuration by Illumina NextSeq 2000. Cell Ranger version 7.1.0 (10x Genomics Inc, CA) was used to map raw data to references of mouse genome mm10-2020-A and Loupe Cell Browser (10x Genomics Inc, CA) to obtain differentially expressed genes between specified cell clusters.

#### scRNAseq Data Analysis

Analyses were performed using Seurat (4.3.0.1) in R studio (RStudio 2023.06.0+421). During quality control, cells with feature counts in the range 1000-6250 and cells displaying less than 5% mitochondrial counts were selected. Following quality control, the dataset underwent pipeline processing that included normalization using VST, multi-dimensional reduction and cell clustering, differential testing framework, and visualization.

### Immunofluorescence Staining and Confocal Microscopy

To evaluate TPRV4 localization and LYVE1+ macrophage distribution, mouse lymphatic vessels were carefully dissected as described above. Vessels were cannulated in calcium-free Krebs buffer prior to fixation with cold 2% paraformaldehyde for 15 minutes at room temperature. After initial fixation, lymphatic vessels were removed from cannulas and transferred into one of the wells of a 24-well plate containing 2% paraformaldehyde for an additional 1-hour fixation at 4°C. Vessels were then permeabilized and blocked for non-specific binding using a PBS solution containing 0.1% triton X-100 and 5% donkey serum for 1 hour at room temperature on a rocking platform. Next, vessels were incubated in primary antibodies (LYVE1 and TRPV4 from Novus Biologicals) at 1:250 in PBS containing 5% donkey serum overnight at 4*°*C. The following day, vessels were washed three times with PBS (total washing time 2 hours) before incubation with secondary antibody (1:500) in PBS containing 5% donkey serum overnight at 4*°*C. Vessels were then washed three times with PBS (total washing time 2 hours) prior to DAPI staining (1:1000) at room temperature for 5 min. Lymphatic vessels were then rinsed once in PBS, and re-cannulated and pressurized for confocal imaging. Immunofluorescence images were collected using a high-resolution and high-speed confocal imaging platform ANDOR Dragonfly 202 (Oxford Instruments), equipped with 4 laser lines (405, 488, 561, and 637 nm), and a Zyla PLUS 4.2 Megapixel sCMOS camera.

### Lymphatic Endothelial Cell Tube (LECT) Isolation

LECTs were isolated from microdissected lymphatic vessels to assess transcript levels of *Trpv4*, prostanoid synthases, and prostanoid receptors. Isolated lymphatic vessels were lightly digested with 1 mg/mL papain, 1 mg/mL collagenase A, 1 mg/mL dispase II, 1 mg/mL trypsin inhibitor, and 1 mM DTT for 25 min at 37°C. Post-digestion, lymphatic endothelial tubes were extracted using a nanoinjector to gently detach and pull the endothelial tube away from the rest of the vessel. LECTs were then placed in an extraction buffer and prepared for RT-PCR.

### Solutions and Chemicals

Krebs buffer, used for both dissection and cannulation, was prepared with 146.9 mM NaCl, 5 mM D-glucose, 4.7 mM KCl, 3 mM NaHCO_3_, 2 mM CaCl_2_·2H_2_O, 1.5 mM Na-HEPES, 1.2 mM MgSO4, 1.2 mM NaH_2_PO_4_·2H_2_O, and 0.5% BSA (pH = 7.4). The solution was then sterile filtered prior to storage at 4°C. A separate Krebs buffer that did not contain BSA was used for continuous perfusion during pressure myography experiments. For dissection of lymphatic vessels for scRNAseq, low-calcium physiological saline solution (PSS) was prepared with the following: 137 mM NaCl, 10 mM HEPES, 10 mM glucose, 5 mM KCl, 1 mM MgCl_2_, and 0.1% BSA (pH = 7.4) and sterile filtered before storage at 4°C.

Pharmacological agents utilized in pressure myography experiments: GSK2193874 (100 nM, GSK219), GSK1016790A (10-100 nM, GSK101), indomethacin (10 µM, Indo), Terutroban (20 nM, S18886), Nicardipine (1 µM), and Caffeine (10 mM).

Chemicals and reagents were purchased from Sigma-Aldrich (St. Louis, MO) and TOCRIS, and these are listed in a Major Resources Table (Supplemental Table 1).

### RT-PCR

Total RNA was extracted from intact inguinal axillary lymphatic vessels (IALVs) and their corresponding lymphatic endothelial cell tubes (LECT) using the Arcturus PicoPure RNA isolation kit (Thermo Fisher Scientific, Waltham, MA) with on-column DNase I treatment (Qiagen, Valencia, CA) according to the manufacturer’s instructions. RNA was eluted with 12 μL extraction buffer. Purified RNA was then used for cDNA synthesis via reverse transcription as per protocol by iScript cDNA Synthesis Kit (Bio-Rad Laboratories, Hercules, CA). All PCR reactions (25 µL) contained first-strand cDNA mixture as the template, PCR Master Mix 2X (Thermo Scientific, Waltham, MA) 0.5 µM primers (IDT, Coralville, IA). The PCR program comprised an initial denaturation step at 95°C for 5 minutes; followed by 36 cycles of denaturation for 30 seconds at 95°C, annealing for 30 seconds at 57°C, and elongation for 30 seconds at 72°C; and a final extension step at 72°C for 5 minutes. The expected size of PCR products and primer sequences are listed in Supplemental Table 2. PCR amplification products were separated on a 2% agarose gel (E-Gel, ThermoScientific) by electrophoresis and the DNA bands were visualized by SYBR Safe imaging.

### Culture of Human Dermal Lymphatic Endothelial Cells (HDLECs) and Assessment of Cell Confluency

Human Dermal Lymphatic Endothelial Cells (HDLECs) were obtained from PromoCell (Heidelberg, Germany) at passage 2 (Catalog #C-12217). Cells were grown in PromoCell Endothelial Cell Media containing Supplement Pack (Catalog # C-22110) and maintained at 37°C and 5% CO_2_. Cells were plated at passage number 3 at 1,000 cells per well and allowed to grow for 24 hours. 12-well replicates were then treated with: 1) Control (1:1000 DMSO), or 2) TRPV4 agonist (10 nM GSK101), or 3) TRPV4 antagonist (100 nM GSK219). All drugs were diluted from stocks in complete growth medium. Cell confluency was immediately measured at 2 minutes and monitored over 48 hours, with measurements taken every 4 hours, in a Sartorius Incucyte SX5 live cell imaging system (Sartorius, Gottingen, Germany). Percent change in confluence at each 4-hour time point was calculated based on the confluence at 2 minutes. Results were plotted as the mean percent change versus time for each treatment and averaged for 3 experiments.

### Statistical Analyses

Results were analyzed with GraphPad Prism Version 10.2.2. Unpaired parametric t-test, two-way ANOVA corrected with Tukey test for multiple comparisons, and one-way ANOVA corrected using Dunnett test were used to determine differences in contractile function parameters, with statistical significance set at p<0.05. Statistical tests were specified in the captions of each figure legend.

## Results

### Pharmacological Activation of TRPV4 Channels Leads to Contractile Dysregulation of Collecting Lymphatic Vessels and Uncovers a Sex Difference

To investigate the functional role of TRPV4 channels in regulating lymphatic contractility, inguinal axillary lymphatic vessels (IALVs) were dissected, cannulated, and prepared for pressure myography experiments as previously described^47^. We first determined the acute effects of pharmacological activation of TRPV4 channels in IALVs isolated from C57BL6/J (WT) male and female mice. The contractile activity of lymphatic vessels was first recorded under control conditions for a period of 2 minutes. Then, vessels were exposed to the potent and selective TRPV4 agonist, GSK1016790A (GSK101, 10 nM), while the pressure myography chamber bath was maintained under constant perfusion with fresh BSA-free Krebs buffer (0.18 mL/min in a 2.5 mL total volume bath). The functional response to pharmacological activation of TRPV4 channels was followed for 10 minutes as GSK101 washed out with perfusion. Interestingly, the response to GSK101 was multiphasic (Figure 1A) and included, an initial robust constriction (i.e., full occlusion of the lymphatic vessel while displaying very high frequency, very low amplitude contractions, referred to as rigor) that was sustained for at least 1 minute and in some instances remained for over 4 minutes. Subsequently, there was a progressive relaxation of the vessel with contractile frequency going from high to low, and contraction width and ejection fraction going from low to high (Figure 1A, label “a”, and Figure 1B-E). Next, there was partial or complete inhibition of contractions (Figure 1A, label “b”). Finally, a partial restoration in contractile activity presumably as a function of GSK101 wash-out (Figure 1A, label “c”). Contractile function was assessed in these areas of interest, which are highlighted in the representative STM examples shown in Figure 1A and 1F for lymphatic vessels from male and female WT mice, respectively.

The initial robust and sustained constriction in response to GSK101 was observed in all male vessels. In the period following the initial constriction, IALVs from male mice displayed varying degrees of loss of contractions; some vessels underwent significant vasodilation and complete cessation of contractions (Figure 1A), while in other vessels, some contractile activity remained with significantly reduced frequency and contraction waves that failed to propagate along the entire lymphatic segment (Supplemental Figure 1A). In contrast to the vessels isolated from male mice, IALVs from female mice showed less severe but still significant contractile dysfunction following stimulation with GSK101. Overall, IALVs from female mice had shorter, more localized constriction events (Figure 1F and Supplemental Figure 1B). The time lymphatic vessels spent in rigor was ∼3-fold greater in male (2.01±0.36 min, n=11) compared to female (0.65±0.15 min, n=14) mice (Figure 2A).

We then utilized newly developed Python-based tools to perform automated tracking of inner and outside diameter from a given region of interest along the lymphatic segment and subsequent automated in-depth quantitative analyses of lymphatic contractility. Consistent with our previous studies^46,47,50–57^, these analyses included the calculation of mean values of contraction amplitude, contraction frequency, ejection fraction (EF), fractional pump flow (FPF), end-diastolic diameter (EDD), end-systolic diameter (ESD), and width of contraction. Following stimulation with GSK101, the pumping capacity of all lymphatic vessels from both male and female mice was significantly impaired, this was evident by the significantly decreased contraction amplitude, EF, and FPF. Frequency was only significantly affected (i.e., decreased) in IALVs from male mice (Figure 2B-E). Notably, the decreased EF and FPF remained significantly reduced after 10 minutes of drug washout (Figure 2D,E). Changes in vessel diameter are normally accompanied by changes in contractile frequency. Lymphatic vessels from both male and female mice displayed an increase in baseline diameter, as evident by the increased EDD and ESD following stimulation with GSK101 (Figure 2F,G, labels “b” and “c”). The mean width of contractions (measured at 50% contraction amplitude) trended to be shorter following the sustained constriction; however, these changes did not reach statistical significance (Figure 2H).

**Figure 1.**
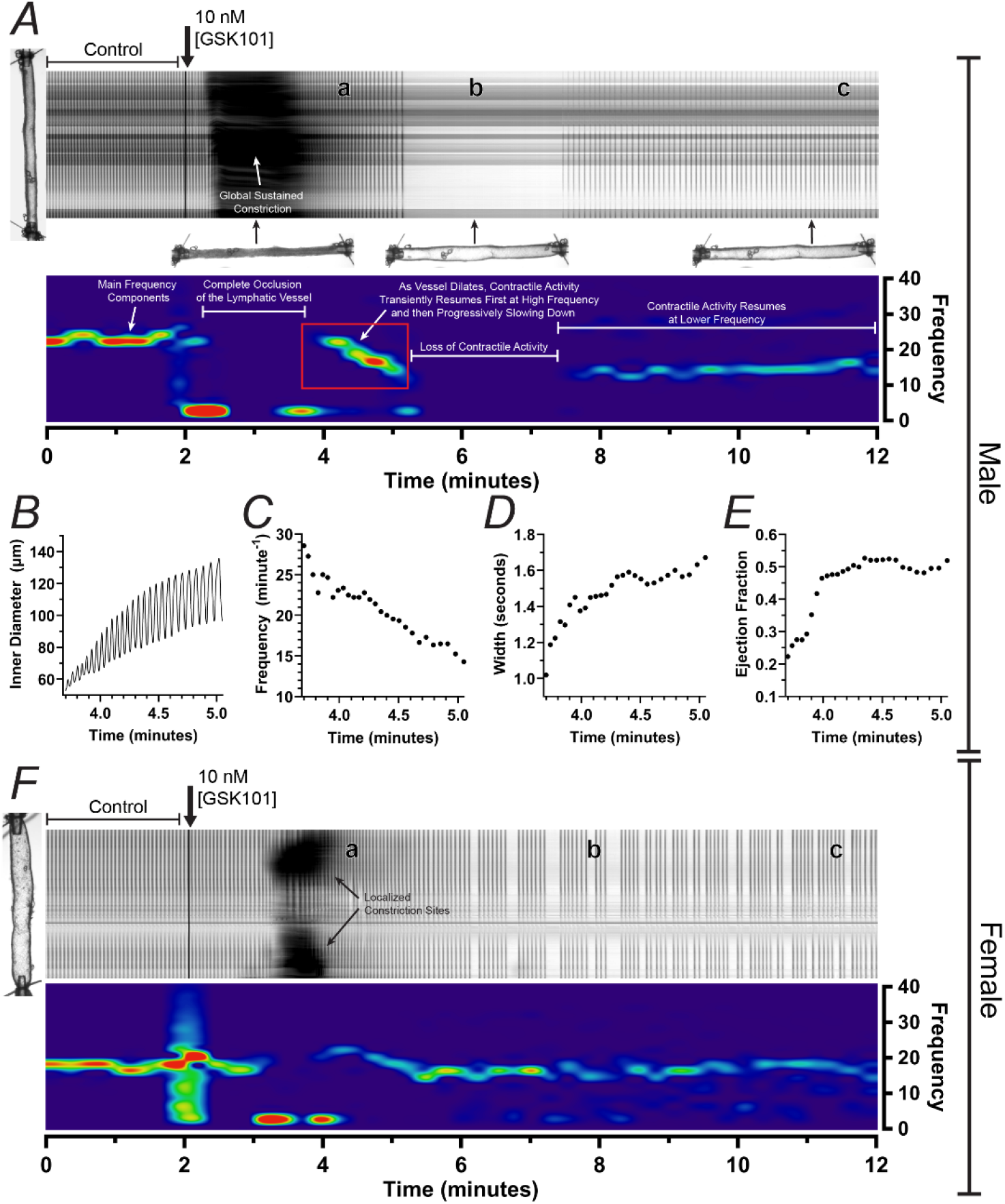
Pharmacological activation of TRPV4 channels results in significant impairment of contractile function of lymphatic vessels in male mice. (A) Space-time map (STM) and corresponding spectrogram generated from a representative 12-minute video recording of the contractile activity of an inguinal axillary lymphatic vessel (IALV) from a male mouse. STM displays the contractility of the vessel under control conditions and upon pharmacological activation of TRPV4 channels using GSK101 (10 nM). Sustained, strong, and global constriction was observed in male IALVs following stimulation with GSK101 (dark area between minutes 2-4), then a transient increase in contractile frequency was observed as rigor ended (label “a” on STM), followed by loss of contractile activity, evident by the lack of narrow dark vertical bands (label “b” on STM), and finally contractile activity is partially restored after 10 minutes of drug washout (label “c” on STM). The spectrogram depicts the main frequency components (major frequency components in bright colors over a dark blue background) as a function of time. (B-E) Contraction-by-contraction analysis for the contractile activity immediately following the sustained constriction (see red box spanning times ∼3.5-5 minutes in the spectrogram). (F) STMs and corresponding spectrogram generated from a representative example of a 12-minute video recording of the contractile activity of the inguinal axillary lymphatic vessel from a female mouse.

**Figure 2.**
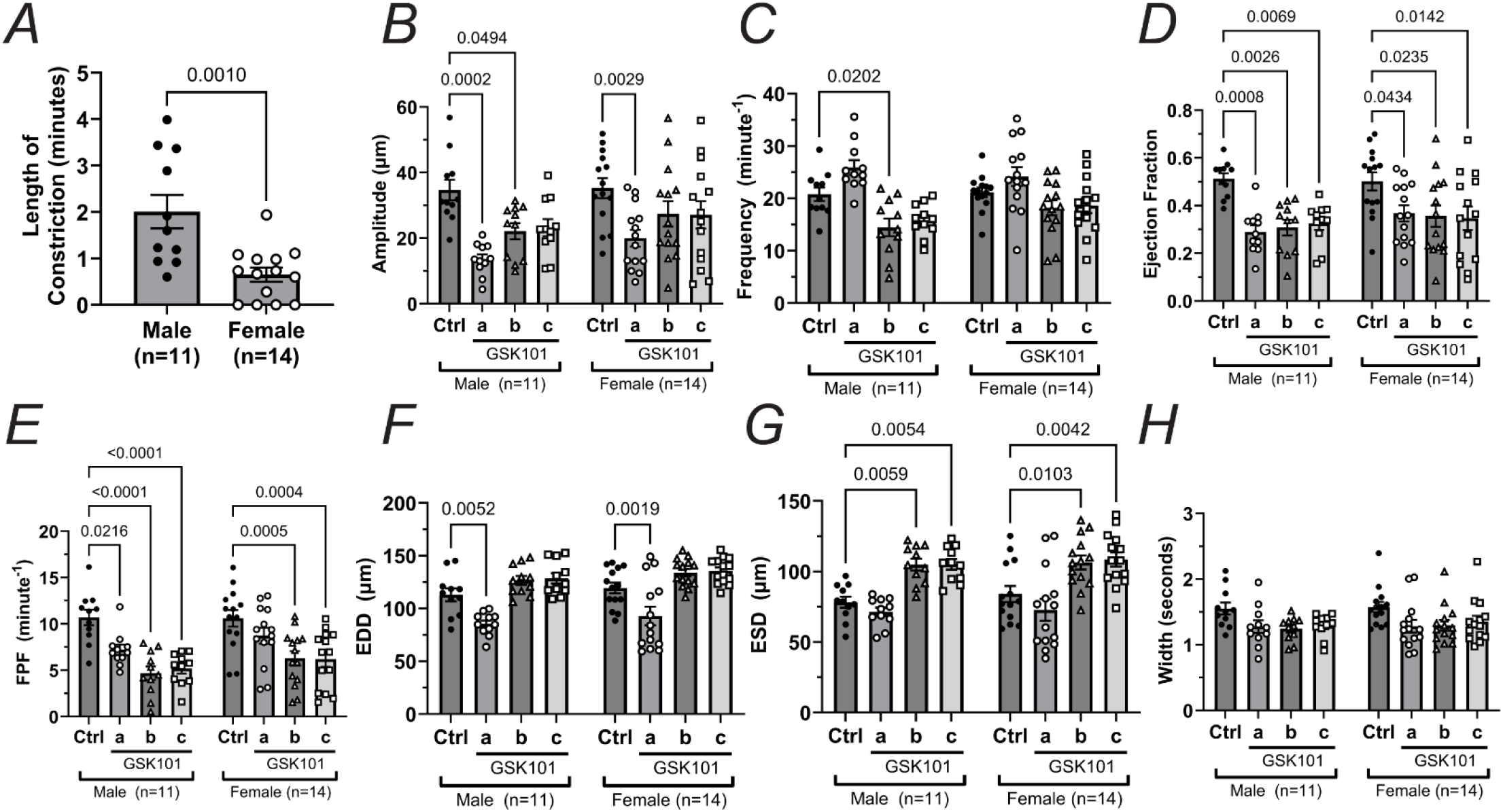
Quantitative assessment of the effects of pharmacological activation of TRPV4 channels on the contractile function of inguinal axillary lymphatic vessels from male and female mice. Summary of the contractile parameters under control conditions and following stimulation with GSK101 (10 nM). Data is separated by sex, with each data point representing an individual lymphatic vessel (n=11 and n=14 for males and females, respectively). **(A)** Length of the constriction induced by GSK101; **(B-H)** Contraction amplitude, frequency, ejection fraction, fractional pump flow (FPF), end-diastolic diameter (EDD), end-systolic diameter (ESD), and contraction width. These parameters were calculated from automated tracking of inner diameter traces. Following stimulation with GSK101, parameters were calculated at 3 different regions depicted with the letters “a-c” during the 10-minute drug washout period and compared with an initial 2-minute period in control conditions. Values are reported as mean±SEM. Statistical significance was determined using an unpaired parametric t-test for data in (A) and two-way ANOVA corrected for multiple comparisons using a Tukey test for data in (B-H). Significant differences are shown for p values <0.05. For simplicity, only significantly different comparisons using the control group, i.e., Ctrl, as reference are shown.

### Lymphatic Vessels and Arterioles from Various Anatomical Regions Display Different Responses to Activation of TRPV4 Channels

To further examine the extent to which TRPV4 channels regulate the function of collecting lymphatic vessels in different anatomical regions, we conducted additional experiments on popliteal afferent, superficial cervical, and mesenteric lymphatic vessels. Similar to IALVs, upon stimulation with GSK101, popliteal and superficial cervical lymphatic vessels also exhibited a significant constriction and subsequent impairment of contractile function; however, constriction was often localized to specific sites and shorter in duration. Mesenteric lymphatics did not display robust constriction, in a few cases, only localized sites of very low amplitude constrictions were observed (Figure 3). Despite the presence or absence of the initial constriction, the popliteal and superficial cervical lymphatics displayed subsequent inhibition of contractions, as evident by the increased space between vertical bands in the representative STM examples shown in Figure 3A,B. Mesenteric lymphatic vessels from the mouse do not consistently develop robust propulsive contractions and therefore, changes in contractile frequency were not assessed (Figure 3C).

We also examined the effects of pharmacological activation of TRPV4 channels in arterioles from the inguinal axillary and mesenteric regions. Importantly, these arterioles displayed similar responses as observed in lymphatics from their corresponding regions, i.e., arterioles from the inguinal axillary region displaying a robust and sustained constriction following stimulation with 10 nM GSK101, while this was not observed in mesenteric arterioles, which only displayed a gain in tone and localized sites of very low amplitude constrictions (Supplemental Figure 2).

**Figure 3.**
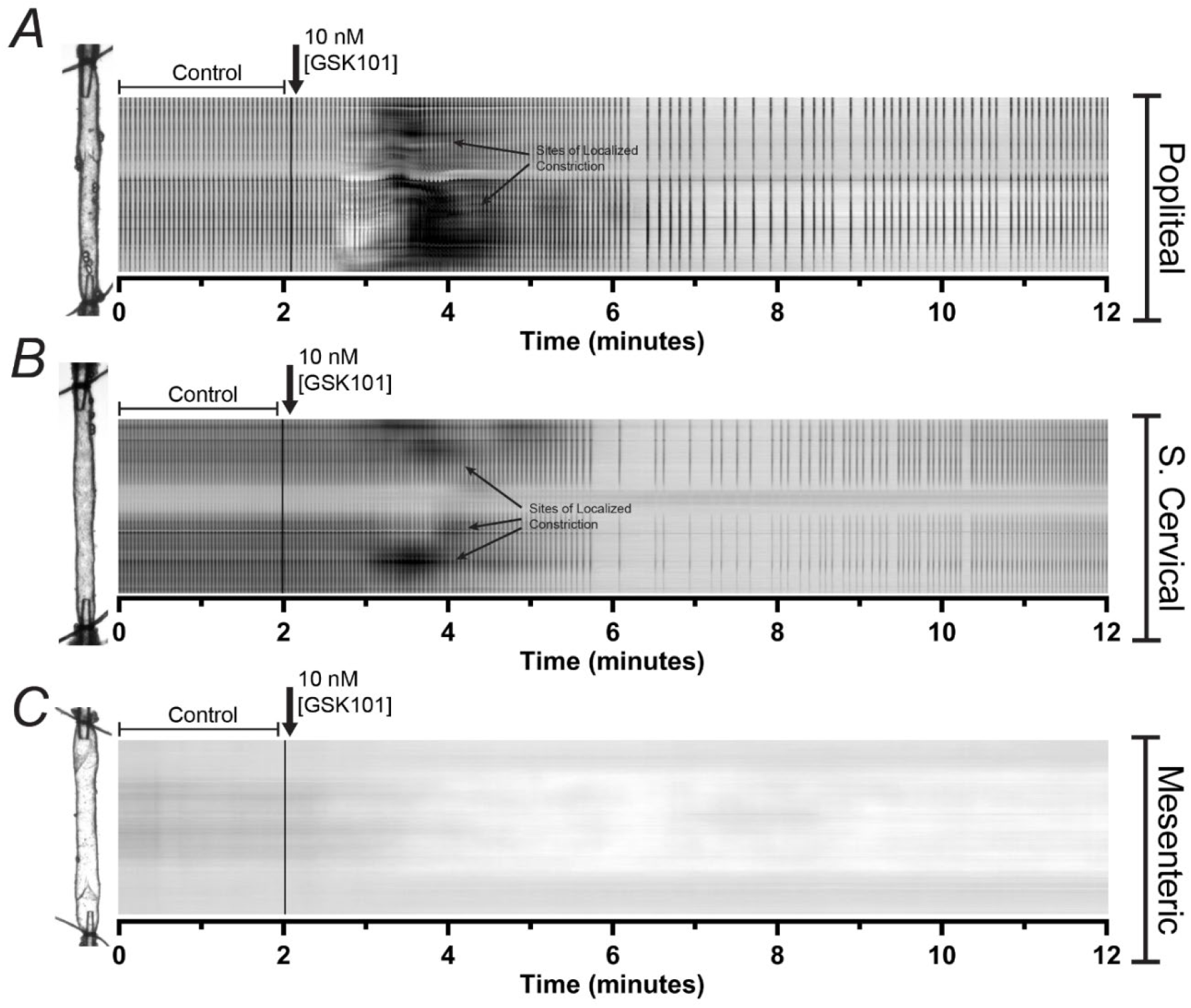
Varying modulation of contractile activity mediated by TRPV4 channel activation is observed across different anatomical regions in mice. Representative STM examples for experiments performed on popliteal afferent, superficial cervical, and mesenteric lymphatic vessels from the mouse. Upon pharmacological activation of TRPV4 channels, both popliteal and superficial cervical lymphatic vessels displayed a sustained and robust constriction similar to that observed in IALVs **(A,B)**, while mesenteric lymphatic vessels did not display any robust constriction **(C)**. Following the initial constriction, all vessels displayed vasodilation, and in the case of popliteal and superficial cervical lymphatics, an impairment in contractile function was consistent with being mediated by nitric oxide.

### Lymphatic Vessels from Different Species Display a Similar TRPV4-Mediated Regulation of Contractile Function

In addition to anatomical site differences, the mechanisms that regulate vascular and lymphatic function may also differ across species. Recent studies using mesenteric or diaphragmatic lymphatic vessels from the rat did not report the presence of a robust constriction associated with activation of TRPV4 channels following flow-induced activation or pharmacological activation (GSK101, 50-350 nM), respectively^35,58^. Therefore, we performed experiments to test lymphatic vessels from additional anatomical regions from rats to determine if the strong vasoconstriction response we observed in the mouse was exclusive to this species. We assessed the functional response to 100 nM GSK101 in rat mesenteric, popliteal afferent, superficial cervical, and inguinal axillary lymphatics. This is the concentration that followed the reported half maximal effective concentration, i.e., EC50, of 31 nM, as determined by the measurement of the increase in intracellular calcium, as reported in previous studies^58^. Our results confirmed that, under no flow conditions, rat mesenteric lymphatic vessels do not display any significant TRPV4-mediated impairment of contractile function, i.e., no detected strong vasoconstriction nor significant decrease in contractile activity (Figure 4A). However, upon stimulation with GSK101, rat popliteal and superficial cervical vessels displayed a strong global constriction and impaired/erratic contractile frequency (Figure 4B,C). Rat inguinal axillary lymphatic vessels showed localized constriction sites (Figure 4D). Finally, we also tested mesenteric lymphatic vessels from non-human primates, i.e., rhesus macaque (monkey). Consistent with mouse and rat mesenteric lymphatic vessels, these also did not show strong vasoconstriction when stimulated with GSK101; however, they displayed reduced contractile frequency (Figure 4E).

**Figure 4.**
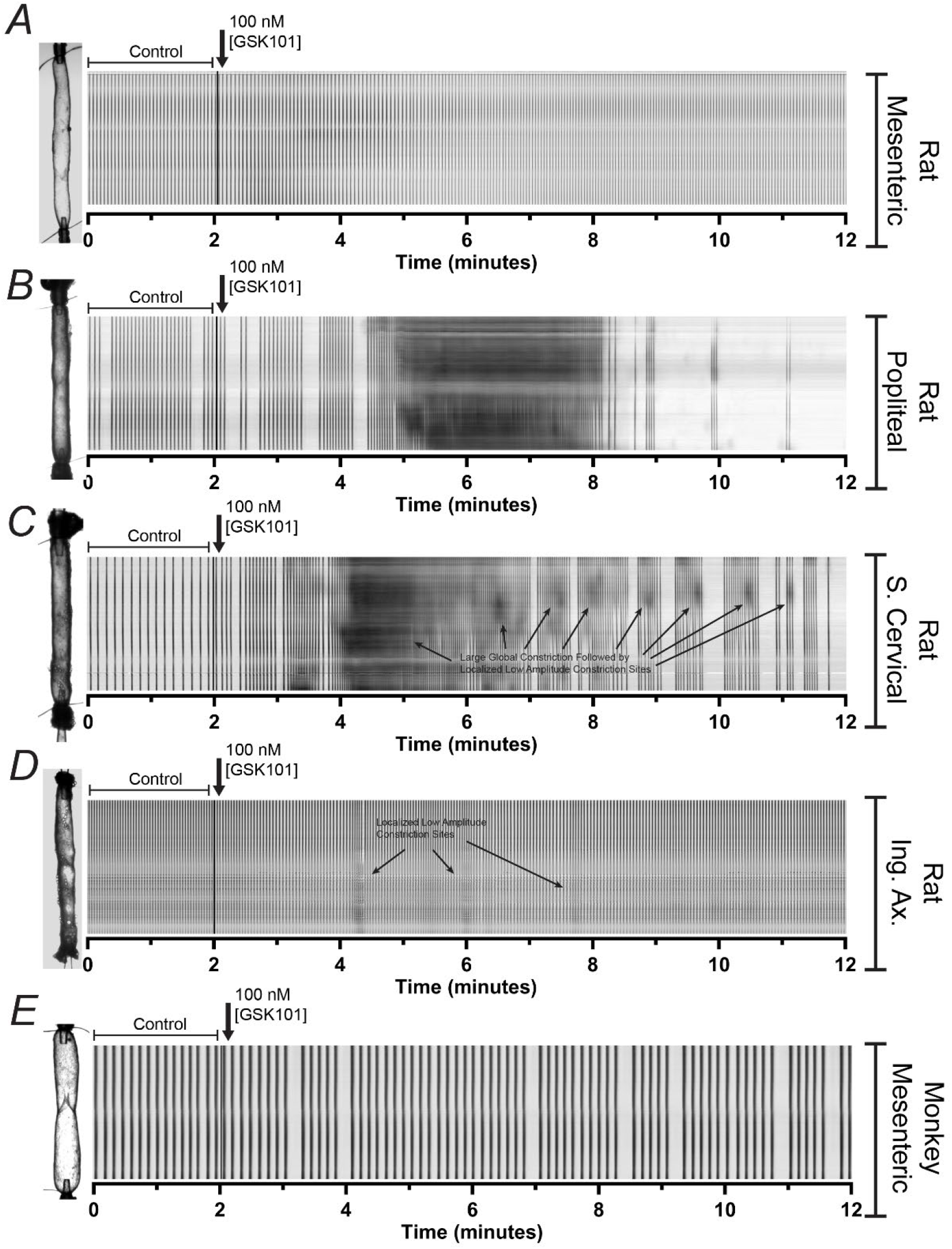
Varying modulation of contractile activity mediated by TRPV4 channel activation is observed across different species. Representative STM examples for experiments performed on **(A)** rat mesenteric, **(B)** rat popliteal afferent, **(C)** rat superficial cervical, **(D)** rat inguinal axillary, and **(E)** rhesus macaque (monkey) mesenteric. Upon stimulation with GSK101 (100 nM), rat popliteal and superficial cervical vessels displayed a strong and sustained constriction followed by significant impairment of contractile activity; however, rat mesenteric lymphatics remain largely unaltered. Following exposure to GSK101, mesenteric lymphatic vessels from rhesus macaque did not display an apparent constriction, but contractility was partially impaired (i.e., decreased frequency). These are representative examples from a total of n=7 experiments.

### TRPV4 Channels are Dispensable for Normal Pacemaking of Healthy Collecting Lymphatic Vessels

As mentioned previously, other studies have implicated TRPV4 channels in the modulation of the intrinsic contractility of collecting lymphatic vessels via temperature and flow sensing^35,58^. To determine the extent to which TRPV4-mediated intracellular calcium activity plays a role in regulating the normal contractility of lymphatic vessels under no flow conditions, we used the TRPV4-channel potent and selective inhibitor, GSK2193874 (GSK219, 100 nM). Following a 20-minute incubation with GSK219, no significant changes were observed in the contractile function of IALVs (Figure 5A), as evident by unaffected contractile parameters, including amplitude, frequency, and EF. There were also no observed changes in EDD nor width of contractions (Figure 5B-F). These results suggest that under no flow conditions, the baseline activity of TRPV4 channels does not play a critical role in regulating the contractile activity of healthy lymphatic vessels and is dispensable for lymphatic muscle cell (LMC) pacemaking. In addition, pharmacological inhibition of TRPV4 channels with GSK219 prevented any impairment of the lymphatic contractile function induced by GSK101 (Figure 5) in mouse IALVs.

**Figure 5.**
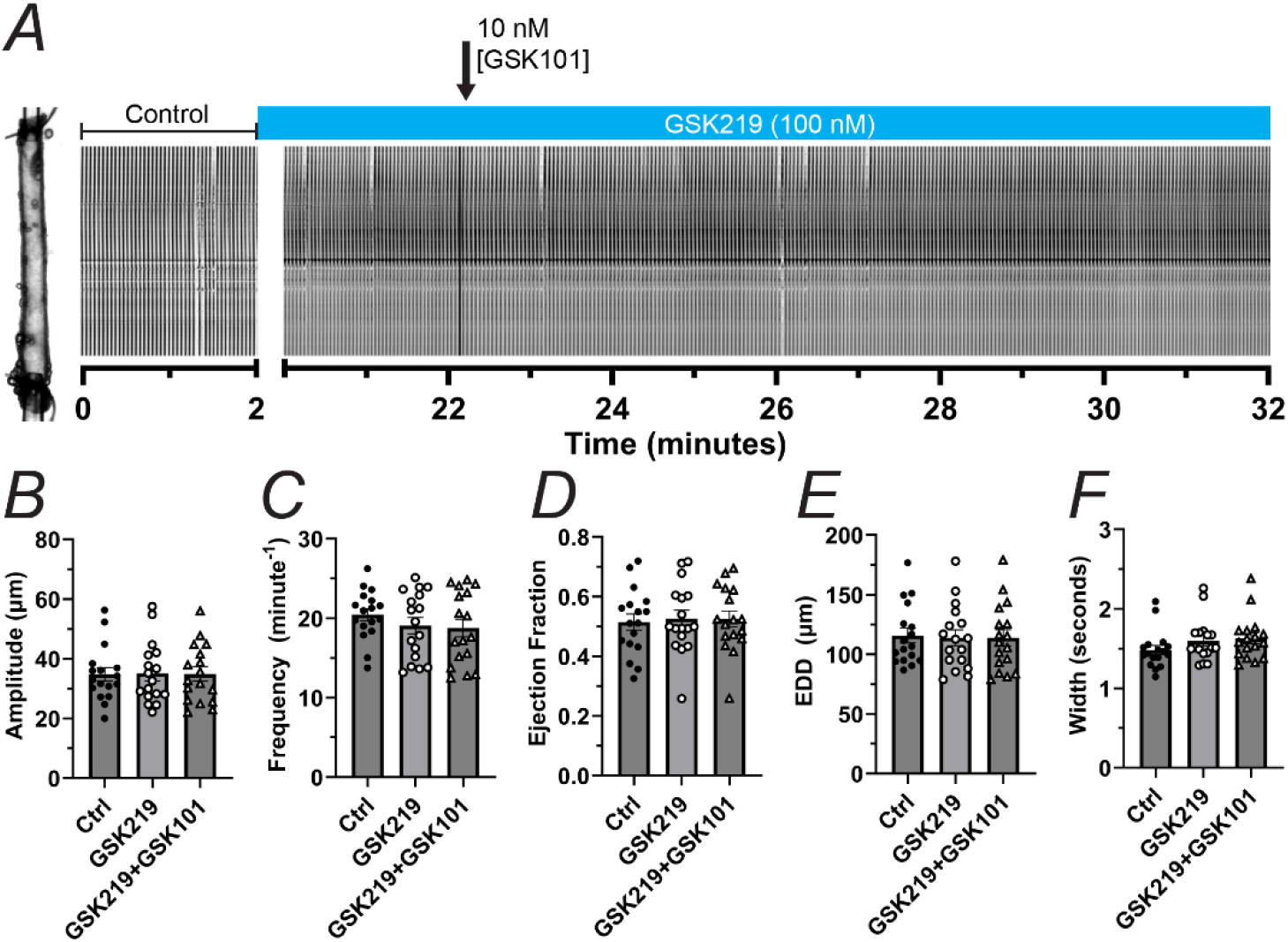
Pharmacological inhibition of TRPV4 channels prevents the functional impairment induced by GSK101. **(A)** Representative STM example depicting the contractile activity of an inguinal axillary vessel from the mouse under control conditions and following stimulation with GSK101 (10 nM) in the presence of TRPV4 antagonist GSK219 (100 nM, 20-minute pre-incubation). **(B-F)** Summary of lymphatic contractile parameters, including contraction amplitude, contraction frequency, ejection fraction, end-diastolic diameter (EDD), and contraction width. Compared to control conditions, no significant differences in contractile function were observed after a 20-minute pre-treatment with GSK219 nor following stimulation of GSK101. A total of n=17 experiments, including male and female, were conducted. The concentration of GSK219 was maintained constant throughout the experimental protocol, as indicated by the blue rectangle above the STM. Values are reported as mean±SEM. Statistical significance was determined using one-way ANOVA corrected for multiple comparisons using a Dunnett test. Significant differences are shown for p values <0.05.

### Activation of TRPV4 Channels Leads to the Release of Nitric Oxide and Prostanoids to Impair Lymphatic Contractility

The previous study by DuToit, el al.^58^ demonstrated that a flow-dependent decrease in pumping of rat mesenteric lymphatics mediated by TRPV4 channels involves the release of endothelium-derived mediators including nitric oxide and vasodilatory prostanoids, as this was prevented by the competitive nitric oxide synthase (NOS) inhibitor L-NAME (100 µM) and the non-selective cyclooxygenase (COX) inhibitor indomethacin (10 µM) respectively. In our experiments using mouse IALVs, the reduction in lymphatic pumping (i.e., vasodilation, and decrease in frequency, EF, and FPF) following the initial, transient constriction, was associated with activation of TRPV4 channels and release of nitric oxide from LECs, as this was prevented in the presence of 100 µM L-NAME (Figure 6A-F). Endothelial TRPV4 channels have also been implicated in promoting lymphangiogenesis and angiogenesis during mouse hindlimb ischemia. Yamada et al. demonstrated that in post-hindlimb ischemia, there was an increase in TRPV4 expression, which promoted the formation of new lymphatic and blood capillaries^59^. We then decided to test the effects of GSK101 in the regulation of cell proliferation in cultures of human dermal lymphatic endothelial cells (HDLECs) followed for a period of 48 hours using an Incucyte S3 system. Cell cultures stimulated with 10 nM GSK101 consistently displayed a significantly increased percent change in cell confluency, while no changes were observed in cultures treated with the TRPV4 antagonist GSK219 when compared to controls (Figure 6G,H).

Our experiments in mouse IALVs suggested that upon activation of TRPV4 channels using GSK101, both vasodilatory and vasoconstrictive factors were being secreted; although, an alternative possibility is that TRPV4 channel expression is present in LMCs, and their expression is heterogenous across anatomical regions and contribute to the vasoconstriction by GSK101. To fully elucidate the mechanism underlying the GSK101-mediated constriction, we explored the potential involvement of vasoconstrictive prostanoids, like thromboxanes. We assessed the contractile function of lymphatic vessels and the effects of TRPV4-channel activation using GSK101 in the presence of the non-selective COX inhibitor indomethacin (10 µM). Indomethacin alone did not alter the contractility of lymphatic vessels; however, it completely prevented the robust constriction induced by GSK101, indicating that upon activation of TRPV4 channels, a vasoconstrictive prostanoid was being released. As anticipated, the subsequent nitric oxide-mediated significant decrease in pumping and significant increase in EDD were still present (Figure 7A-F). These observations indicate that modulation of contractile function by TRPV4 channels is mediated by endothelial nitric oxide and other paracrine signaling from cells embedded within the vessel wall and not by direct activation of TRPV4 channels in LMCs.

**Figure 6.**
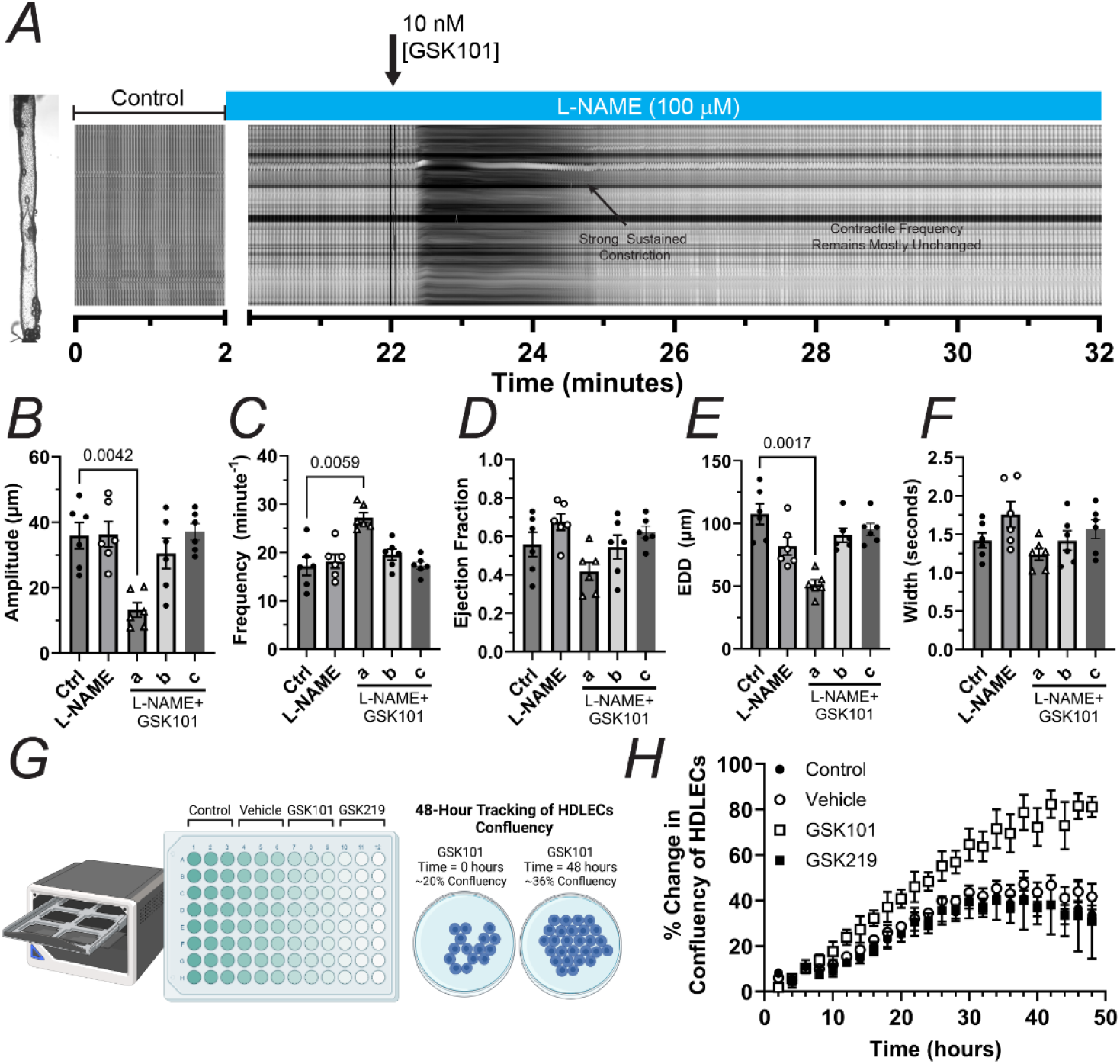
Functional assessment of endothelial TRPV4 channels in whole IALVs and in vitro in cultures of HDLECs. **(A)** Representative STM example depicting the contractile activity of an inguinal, axillary vessel from the mouse under control conditions and following stimulation with GSK101 (10 nM) in the presence of the NOS inhibitor L-NAME (100 µM, 20-minute pre-incubation). **(B-F)** The summary of lymphatic contractile parameters includes contraction amplitude, contraction frequency, ejection fraction, end-diastolic diameter (EDD), and contraction width. Compared to control conditions, no significant differences in contractile function were observed after a 20-minute pre-treatment with L-NAME; however, while L-NAME did not prevent the constriction induced by GSK101, it prevented the subsequent decrease in contractile frequency and pumping capacity. A total of n=6 experiments, including male and female, were conducted. The concentration of L-NAME was maintained constant throughout the experimental protocol, as indicated by the blue rectangle above the STM. **(G-H)** Assessment of the effects of pharmacological activation and inhibition of TRPV4 channels on the proliferation (expressed as percent change in confluency) of human dermal LECs (HDLECs). A total of n=3 experiments were performed each including 12 replicates per experiment. Values are reported as mean±SEM. Statistical significance was determined using one-way ANOVA corrected for multiple comparisons using a Dunnett test. Significant differences are shown for p values <0.05.

**Figure 7.**
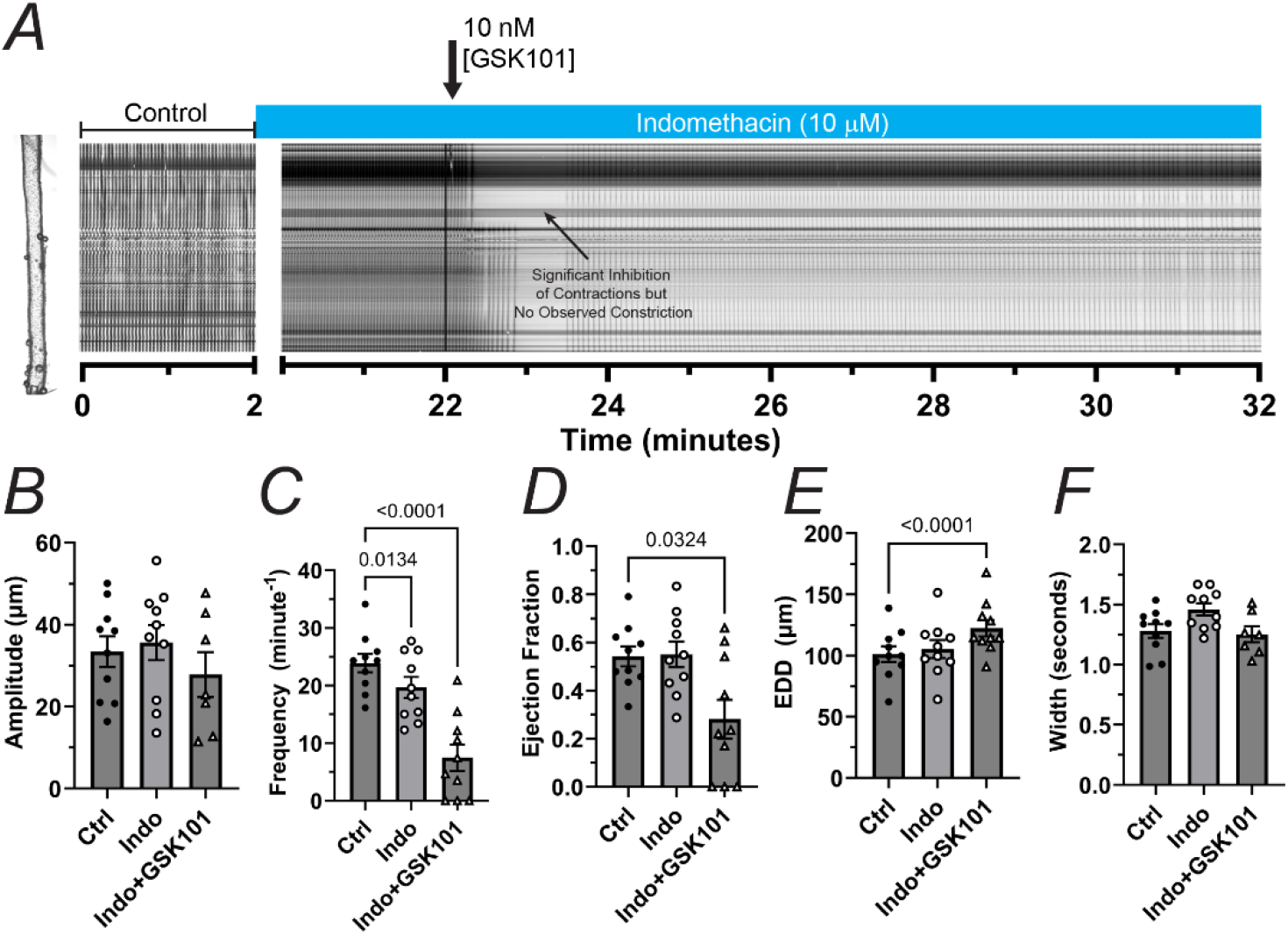
Pharmacological inhibition of cyclooxygenases prevents the functional impairment induced by GSK101. **(A)** Representative STM example depicting the contractile activity of an inguinal axillary vessel from the mouse under control conditions and following stimulation with GSK101 (10 nM) in the presence of the cyclooxygenases COX-1,2 inhibitor (10 µM, 20-minute pre-incubation). **(B-F)** Summary of lymphatic contractile parameters, including contraction amplitude, contraction frequency, ejection fraction, end-diastolic diameter (EDD), and contraction width. A 20-minute pre-treatment with indomethacin prevented the initial robust and sustained constriction induced by GSK101; however, significant impairment of contractile function consistent with that mediated by nitric oxide remained. A total of n=10 experiments, including male and female, were conducted. The concentration of indomethacin was maintained constant throughout the experimental protocol, as indicated by the blue rectangle above the STM. Values are reported as mean±SEM. Statistical significance was determined using one-way ANOVA corrected for multiple comparisons using a Dunnett test. Significant differences are shown for p values <0.05.

### Single-Cell RNA Sequencing on Isolated Collecting Lymphatic Vessels Identifies a Subset of Macrophages Expressing TRPV4 Channels and Links them to the Release of TXA2

To assess the expression of TRPV4 channels in the different cell types within and around the wall of collecting lymphatic vessels, we performed single-cell RNA sequencing (scRNAseq) using collecting lymphatic vessels from male and female WT mice (n=4 mice per group). Previous studies have generated valuable datasets; however, common limitations encountered when single-cell suspensions are prepared from larger tissues are the low percent of cells of lymphatic origin and the need for red blood cell lysis, which may limit the possibilities for downstream analyses. To maximize the capabilities of this technology and maximize the enrichment of the different lymphatic cell types, we prepared single-cell suspensions from individually micro-dissected lymphatic vessels. Given the small size of these lymphatic vessels, collecting lymphatic vessels from 4 different anatomical regions of the same animal (i.e., inguinal axillary (2 vessels), superficial cervical (4 vessels), popliteal afferent (4 vessels), and mesenteric (4 vessels)) were pooled into a single sample. Following automated clustering, a total of 24 different cell-type clusters (i.e., 0-23) were identified (Figure 8A). Upon determination of cell identities, multiple clusters were associated with macrophages, fibroblasts, and LMCs. This may be explained, at least in part, by the differences in the regional origin of the cells included in this study. 13 different cell types were identified, while the identity of the cells in the last 2 clusters (i.e., clusters 22 and 23) remained undetermined (Figure 8B). A table containing the total number of cells per sample and per cell identity, as well as cell viability are included in Supplemental Tables 3 and 4 (where M1-4 and F1-4 identify the different samples/animals and their corresponding sex, M=male and F=female).

Macrophages displayed the highest expression of *Trpv4*, with Clusters 0 and 6 having the largest number of cells expressing this gene (Figure 8C). *Trpv4* expression was also detected in LECs, mesothelial cells, keratinocytes, blood capillary ECs (cap-ECs), and proliferating cells. Importantly, no significant expression of *Trpv4* was detected in LMCs. Prostanoids are active lipid mediators that originate from free essential fatty acids that get catalyzed and synthesized by COXs and specific synthases. Cyclooxigenases COX-1 and COX-2 are encoded by *Ptgs1* and *Ptgs2,* respectively, while prostaglandins, prostacyclins, and thomboxanes are encoded by *Ptges*, *Ptges2*, *Ptges3*, *Ptgds*, *Prxl2b*, *Ptgis*, and *Tbxas1*. Our results showed that *Ptges3* is the most abundant synthase across all cell types. Intriguingly, in addition to *Trpv4,* many macrophages also robustly expressed the synthase*, Tbxas1*, which is the enzyme responsible for the production of the potent vasoconstrictor thromboxane A2 (TXA2) (Figure 8C). These results suggested that TXA2 may be responsible for the strong and sustained constriction of LMCs induced upon activation of TRPV4 channels in macrophages. Expression of *Tbxas1* was also observed in dendritic cells (DCs) and proliferating cells; however, the expression of *Trpv4* in these clusters was significantly lower than that in macrophages. Further supporting information was obtained after assessing the expression of the different prostaglandin receptors. We confirmed that TXA2 receptors (TXA2Rs, encoded by *TXA2r*) display the highest expression in LMCs (Figure 8D). Interestingly, LECs, pericytes, and mesothelial cells also showed significant expression of TXA2r. Suggesting that the release of TXA2 from macrophages would not only induce the constriction of LMCs, but it would also activate TXA2Rs in other cell types, including LECs and pericytes, both known to be important players in the regulation of vascular function.

To further validate our scRNAseq data, we first performed RT-PCR on whole IALVs and intact lymphatic endothelial cell tubes (LECTs) isolated as previously described^34,60^. Our results further demonstrated that *Tbxas1* was detected in whole vessels but not in LECTs (Figure 8E-H). Then, to test the involvement of TXA2 in the GSK101-induced constriction, we performed functional experiments on isolated IALVs exposed to GSK101 in the presence of Terutroban (S18886, 20 nM), a potent TXA2R antagonist. Similar to our results using indomethacin, S18886 completely prevented the strong vasoconstriction following the activation of TRPV4 channels (Figure 9A), confirming that TXA2 is responsible for TRPV4-mediated constriction. These vessels still displayed a significant decrease in pumping (i.e., lower frequency and ejection fraction), consistent with nitric oxide-mediated inhibition of contractions (Figure 9B-F).

**Figure 8.**
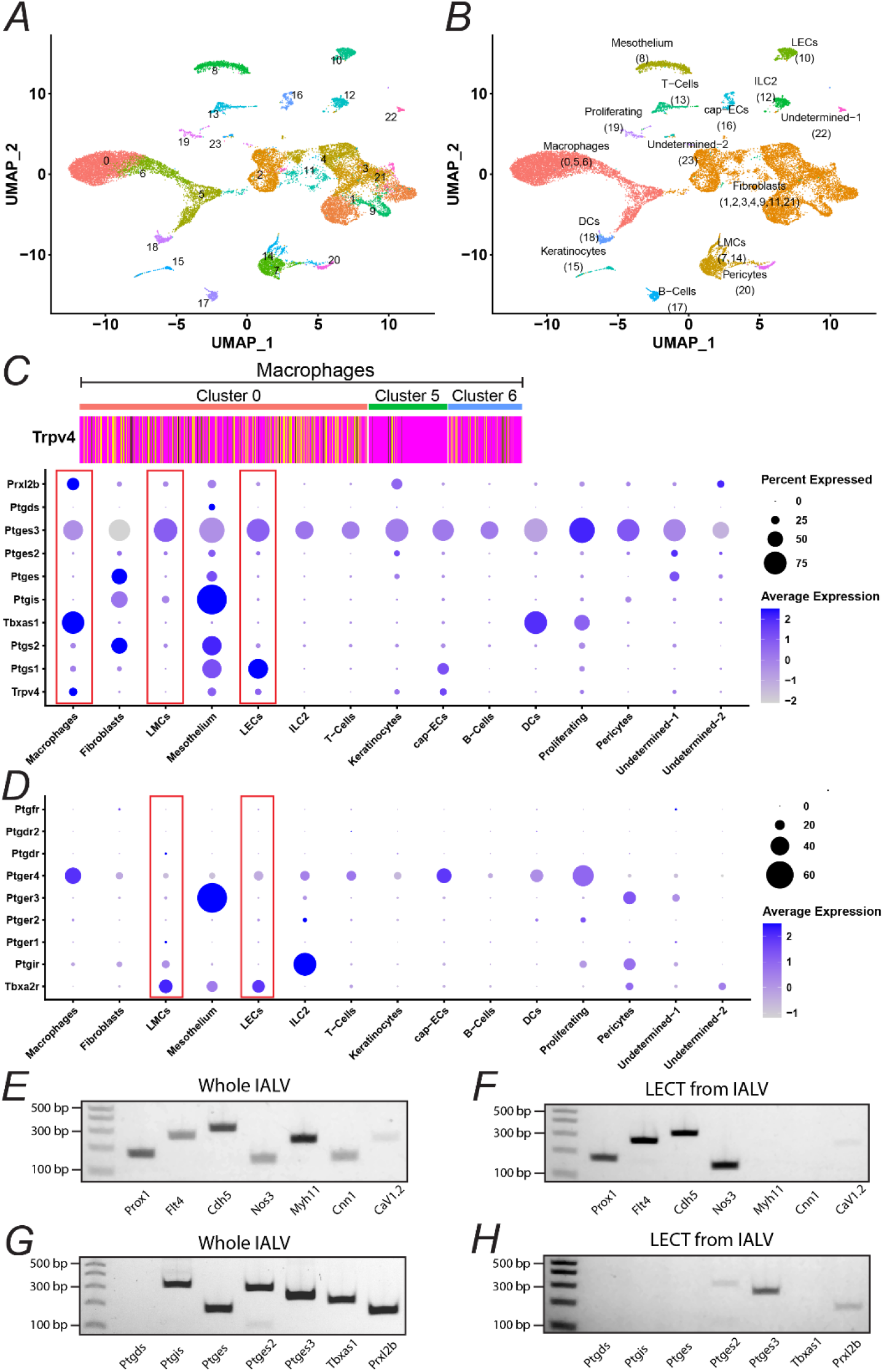
Assessment of expression of *Trpv4* and prostanoid-related genes using single-cell RNA sequencing. Single-cell RNA sequencing (scRNAseq) was completed on samples using isolated lymphatic vessels from 4 different anatomical regions (i.e., inguinal axillary, popliteal afferent, mesenteric, and superficial cervical) from male and female WT mice. **(A)** Differential gene expression analysis identified the 24 distinct cell clusters as shown in this UMAP; **(B)** UMAP with annotated cell types and cluster numbers. Note upon cell identification, multiple clusters were associated with macrophages, fibroblasts, and LMCs. **(C)** DotPlot demonstrating the expression of genes encoding TRPV4, cyclooxygenases, and prostaglandin, prostacyclin, and thromboxane synthases across all different cell types. Gene expressions in macrophages, LECs, and LMCs are highlighted (red boxes). *Trpv4* expression was significantly higher in the macrophage clusters, primarily in clusters 0 and 6, as shown in the upper-insert plot in panel C. **(D)** DotPlot demonstrating expression of the different prostaglandin, prostacyclin, and thromboxane receptors across cell types. Expression in LMCs and LECs are highlighted (red boxes). In these cell types, *TXA2r* encoding the TXA2 receptors display the highest expression in both LMCs and LECs. In the DotPlots in C and D, the percent of cells expressing a gene of interest within a given class is encoded in the size of the dot, while the color encodes the average expression level (expressed as *avg_log2FC*) across all cells within a class. **(E-H)** Representative RT-PCR gels comparing the expression of a 7-gene panel of known markers found in LECs and/or LMCs and a 7-gene panel of all prostanoid synthases in whole IALVs and in isolated lymphatic endothelial cell tubes (LECTs).

**Figure 9.**
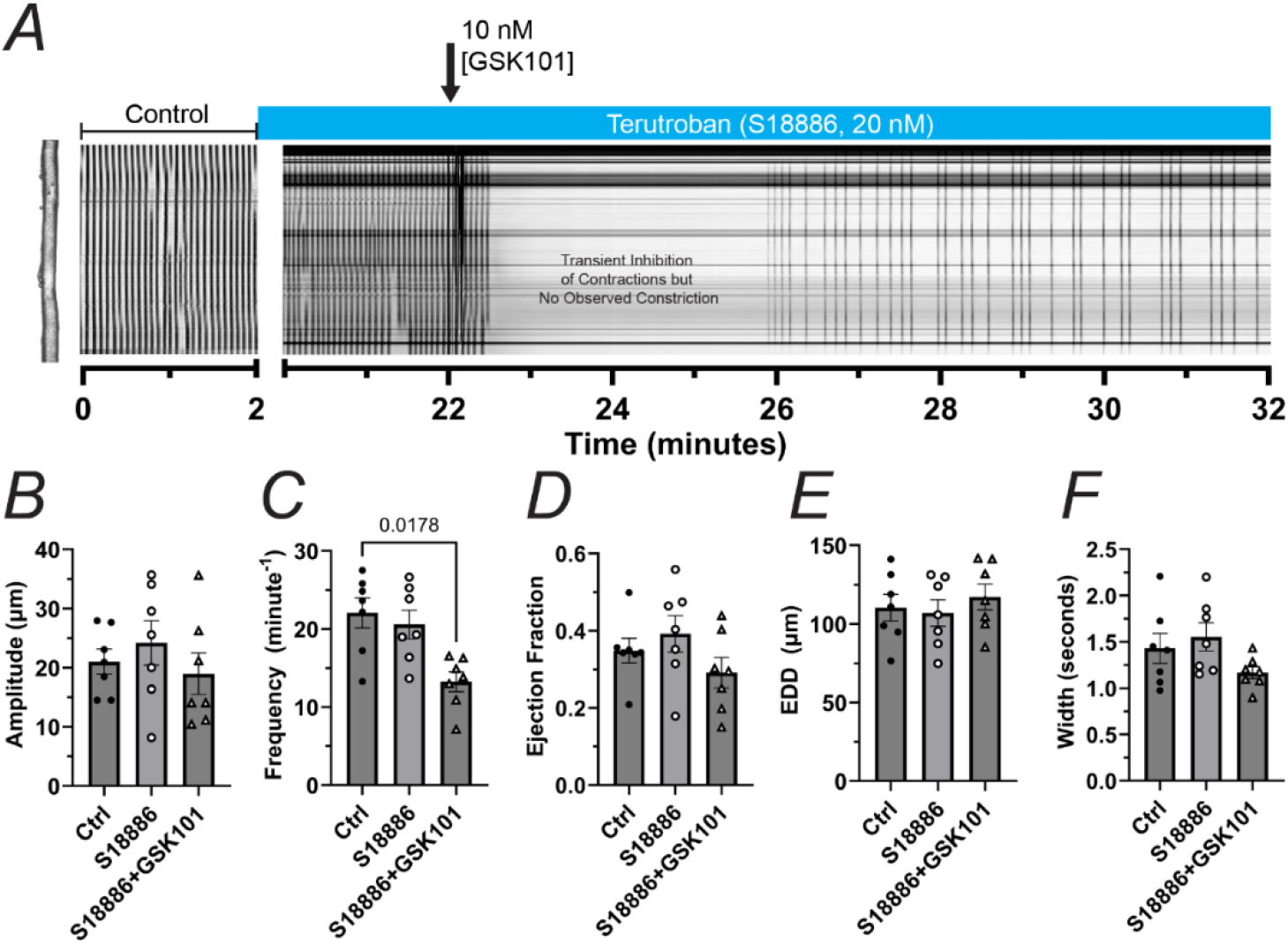
Pharmacological inhibition of TXA2Rs prevents the functional impairment induced by GSK101. **(A)** Representative STM example depicting the contractile activity of an inguinal axillary vessel from the mouse under control conditions and following stimulation with GSK101 (10 nM) in the presence of the potent TXA2R antagonist, terutroban (S18886 20 nM, 20-minute pre-incubation). **(B-F)** The summary of lymphatic contractile parameters includes contraction amplitude, contraction frequency, ejection fraction, end-diastolic diameter (EDD), and contraction width. A 20-minute pre-treatment with terutroban prevented the initial robust and sustained constriction induced by GSK101; however, similar to the results obtained in the presence of indomethacin, significant impairment of contractile function consistent with that mediated by nitric oxide remained. A total of n=6 experiments, including male and female, were conducted. The concentration of terutroban was maintained constant throughout the experimental protocol, as indicated by the blue rectangle above the STM. Values are reported as mean±SEM. Statistical significance was determined using one-way ANOVA corrected for multiple comparisons using a Dunnett test. Significant differences are shown for p values <0.05.

### Trpv4-Expressing Macrophages Differentially Express a Tissue-Resident Macrophage Signature

Utilizing our scRNAseq dataset, we further characterized the subset of macrophages that expressed *Trpv4*. Differential gene expression analysis led to a list of 284 genes significantly different in *Trpv4*-expressing macrophages compared to non-*Trpv4*-expressing macrophages. This list included 86 upregulated genes and 198 downregulated genes. The heatmap shown in Figure 10A includes the top 25 upregulated and downregulated genes based on their *avg_log2FC* expression values. In addition to *Trpv4*, this subset of macrophages also differentially expressed known markers of tissue-resident macrophages (TRMs), including *Cd163*, *Folr2*, *Mrc1*, *Ccl8*, *Apoe*, *Cd209f*, *Cd209d*, *Cd209g,* and the known non-specific lymphatic marker, *Lyve1* (Figure 10A). Although not differentially expressed, expression of *Timd4*, another established marker of TRMs, was also observed in these macrophages. Interestingly, *Ilb1* was the most significantly downregulated gene in *Trpv4*-expressing macrophages. The protein encoded by *Ilb1* is Interleukin 1 Beta (Il-1b), a cytokine known to be produced by macrophages upon activation. Il-1b has been implicated as a major mediator of inflammation. To validate our scRNAseq findings, we performed immunofluorescence staining for TRPV4 and LYVE1 on isolated and pressurized IALVs. Following confocal microscopy imaging, our images confirmed positive and strong staining for TRPV4 channels that colocalized with LYVE1 staining (Figure 10B-G).

**Figure 10.**
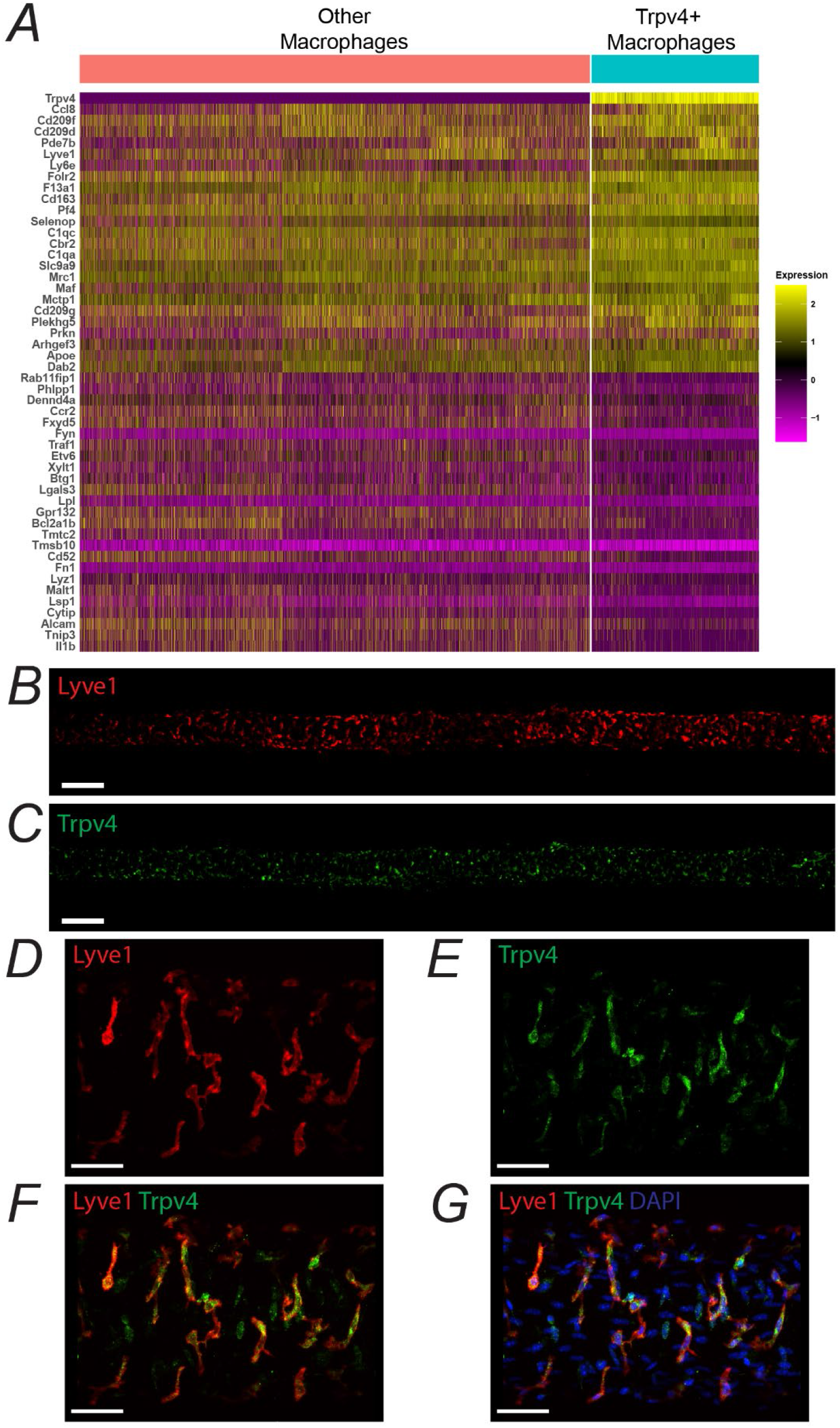
Characterization of *Trpv4*-expressing macrophages via differential gene expression analysis. Macrophages expressing or not expressing *TRPV4* were subclustered, and their genetic make-up was compared by means of differential gene expression analysis. The list of differentially expressed genes was sorted by average expression (i.e., *avg_log2FC*) from highest to lowest values, and the top and bottom 25 genes are reported in the HeatMap in **(A)**; and **(B-G)** Maximum projections from Z-stacks collected using confocal microscopy in isolated and pressure-fixed IALVs stained for TRPV4 (green), LYVE1 (red), and DAPI (blue). Scale bars are 200 µm for panels B-C (10X magnification) and 50 µm for panels D-G (40X magnification).

### Decreased Density of Trpv4-Expressing Macrophages in Female Mice may be Responsible for Sex-Based Differences

Activation of TRPV4 channels resulted in significant and persistent impairment of the contractile function of lymphatic vessels from both male and female mice; however, lymphatic vessels from female mice displayed an apparent partial protection against the detrimental effects of activation of TRPV4 channels in macrophages, resulting in significantly shorter and weaker initial constriction when compared to vessels from male mice (Figures 1 and 2). By using our scRNAseq dataset, we sought to determine if these sex-differences could be explained by differential expression of critically involved genes. We generated subsets of *Trpv4*-expressing macrophages from female and male mice, respectively, and then determined differentially expressed genes. Based on their *avg_log2FC* expression values, the top 25 upregulated and downregulated genes are shown in Figure 11A. *Trpv4*-expressing macrophages from male mice displayed significantly higher expression of the chemokine encoding gene *Ccl8* and the forkhead family transcription factor *Foxo3*; while other chemokine encoding genes *Ccl4* and *Ccl3* were higher in the female group. We then determined and compared the expression of *Trpv4* in *Trpv4*-expressing macrophages, LECs, and LMCs. The expression of *Trpv4* was significantly higher, >8-fold and >80-fold, in *Trpv4*-macrophages when compared to expression in LECs and LMCs, respectively. No significant differences in expression were observed across sexes (Figure 11B). The synthesis and release of the prostanoid TXA2 by macrophages requires the corresponding functional synthase, i.e., *Tbxas1*. No significant differences were found in the expression of *Tbxas1* between sexes either (Figure 11C). We then evaluated whether the sex difference could be explained by the expression of TXA2Rs (encoded by *TXA2Rs*) in LMCs; however, no differences were detected (Figure 11D). When determining the total number of cells per cell type population and per sample, we noticed that samples from female mice contained a significantly lower number of cells, i.e., 2447±149 average total number of cells for female samples versus 2930±25 for male samples (Figure 11E). This was associated with a noticeably smaller size of lymphatic vessels observed during dissection and the lower body weight commonly observed in female mice (21.4±0.6 g) versus that of male mice (28.1±0.8 g) (Figure 11F). Importantly, the percent of total macrophages and the percent of *Trpv4*-expressing macrophages when referenced to the total number of cells, or to the overall macrophage population, were all significantly decreased in samples from female mice (Figure 11G-I). Suggesting that the significantly shorter and weaker constriction induced upon activation of TRPV4 channels in lymphatic vessels from female mice may be explained by a significantly decreased density of *Trpv4*-expressing macrophages when compared to males.

**Figure 11.**
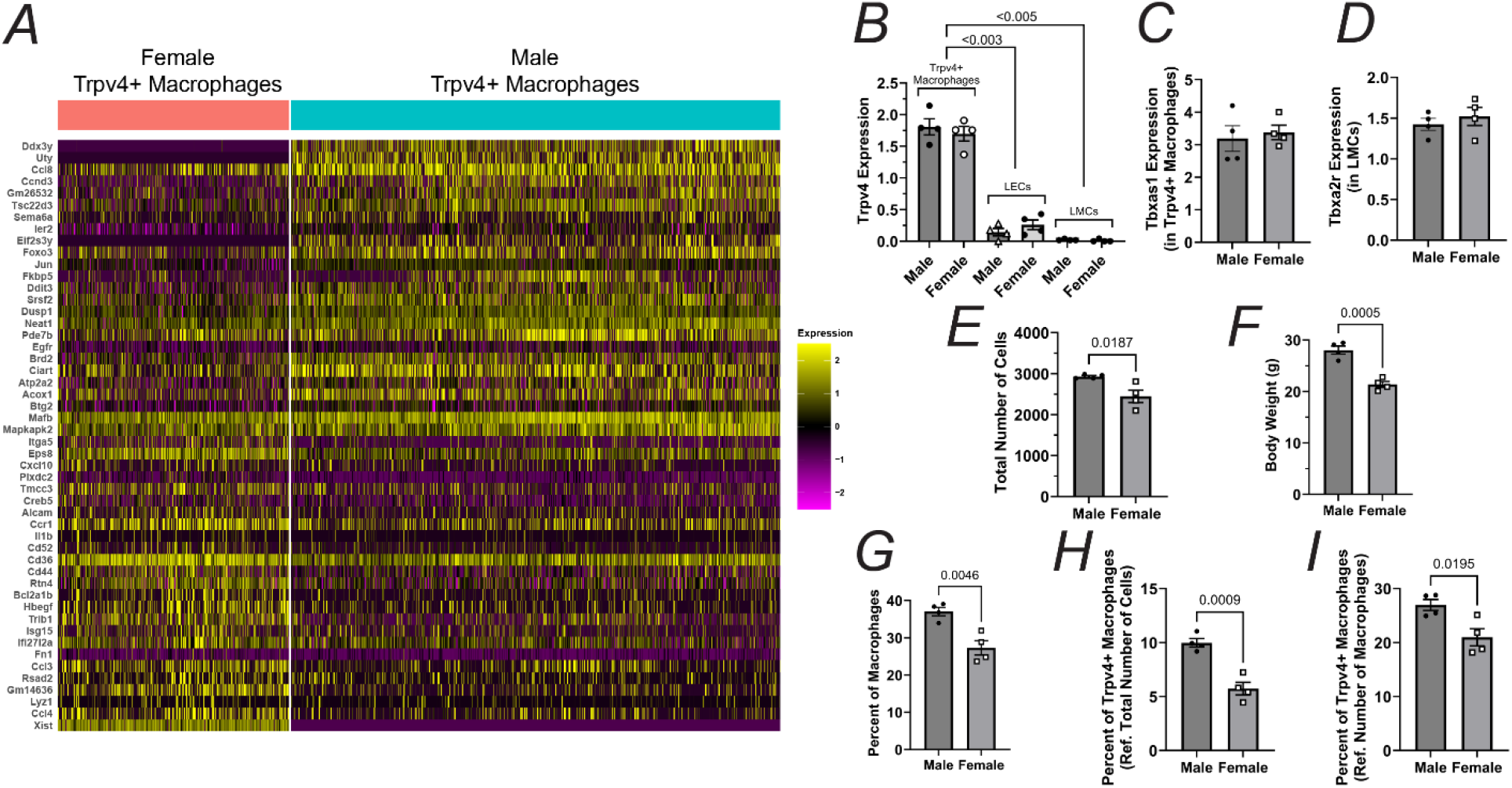
Sex-based comparison of *Trpv4*-expressing macrophages using differential gene expression analysis. Macrophages expressing *TRPV4* were subset by sex and their genetic make-up were compared by means of differential gene expression analysis. The list of differentially expressed genes was sorted by average expression (i.e., *avg_log2FC*) from highest to lowest values, and the top and bottom 20 genes are reported in the HeatMap in **(A)**; **(B)** average expression of *TRPV4* in Trpv4-expressing macrophages, LECs, and LMCs; **(C)** average expression of *Tbxas1* in Trpv4-expressing macrophages; **(D)** average expression of *TXA2r* in LMCs; **(E)** total number of cells per sample (n=4 animals for each sex are included); **(F)** body weight per animal; **(G)** percent of macrophages per sample; **(H)** percent of Trpv4-expressing macrophages (referenced to the total number of cells); and **(I)** percent of Trpv4-expressing macrophages (referenced to the total number of macrophages).

### Macrophage TRPV4-Initiated Constriction of LMCs is Mediated by TXA2Rs and Requires Release of Calcium from Intracellular Stores via Ip3-Receptors

To further elucidate the downstream mechanism following activation of the TXA2Rs by TXA2 leading to a sustained (sometimes lasting ≥4 minutes) and robust constriction, first, we investigated the potential involvement of L-type calcium channels (LTCCs). Similar to previous protocols, we utilized IALVs from the mouse, following equilibration, vessels were exposed to the TRPV4 agonist GSK101 (10 nM) in the presence of the LTCC inhibitor nicardipine (1 µM). As anticipated, nicardipine completely inhibited the contractile function of lymphatic vessels^47^; however, it did not prevent the robust constriction induced by GSK101, suggesting that calcium entry through LTCCs is dispensable for the TXA2-induced constriction (Figure 12A). We then tested whether mobilization of calcium from intracellular stores via ryanodine receptors (RyRs) or inositol triphosphate receptors (Ip3Rs) was required. A set of vessels were stimulated with GSK101 following treatment with 10 mM caffeine, which depleted all calcium from intracellular stores via activation of RyRs. As previously reported^61^, caffeine induced transient and strong constriction of the lymphatic vessels, followed by a complete inhibition of spontaneous contractions. Subsequent stimulation with GSK101 did not induce sustained and robust constriction (Figure 12B). Next, we assessed the effects of GSK101 on the contractile function of lymphatic vessels from *Myh11*-CreER^T2^;*Ip3r1*^fx/fx^;*Ip3r2*^fx/fx^;*Ip3r3*^fx/fx^ mice, lacking all three isoforms of Ip3Rs in smooth muscle cells. Lymphatic vessels from *Myh11*-CreER^T2^;*Ip3r1*^fx/fx^;*Ip3r2*^fx/fx^;*Ip3r3*^fx/fx^ mice displayed a significantly lower contractile frequency than those from WT controls, this is in association with the critical role that Ip3r1 plays in the regulation of pressure-dependent chronotropy and contractility, as recently shown by our groups^57,62,63^. Upon stimulation with GSK101, lymphatic vessels lacking Ip3Rs in LMCs did not display robust and sustained constriction (Figure 12C). No significant changes were observed in contraction amplitude, ejection fraction, or EDD. Frequency had a transient increase; however, baseline contractility was restored within 1-2 minutes (Figure 12D-G). These results indicate that upon activation of TXA2Rs, calcium mobilization from intracellular stores into the cytosol is mediated by Ip3Rs and likely leads to subsequent calcium entry from the extracellular space, i.e., store-operated calcium entry (SOCE). Finally, we evaluated the expression of genes encoding channels known to play a part in SOCE. Figure 13 summarizes the expression of *Stim1,2*, *Orai1-3*, *Cracr2a-b*, and *Trpc1,3-7*. Importantly, in LMCs, expression for both *Stim1* and *Stim2* was detected, although strong dominance was observed by *Stim1*; expression of *Orai1* and *Orai3* were significant, with *Orai3* being the highest; no significant expression of *Cracr2a* and *Cracr2b* was detected; and finally, expression of *Trpc1*, *Trpc4*, and *Trpc6* was detected, although *Trpc1* and *Trpc6* were strong dominants. Altogether, the results presented here indicate that upon activation of TRPV4 channels in LYVE1-expressing macrophages, TXA2 is secreted and binds to TXA2Rs in LMCs to induce an increase in intracellular calcium via IP3Rs and promotes SOCE leading to sustained vasoconstriction of lymphatic vessels prior to release of nitric oxide from LECs which promotes vasodilation.

**Figure 12.**
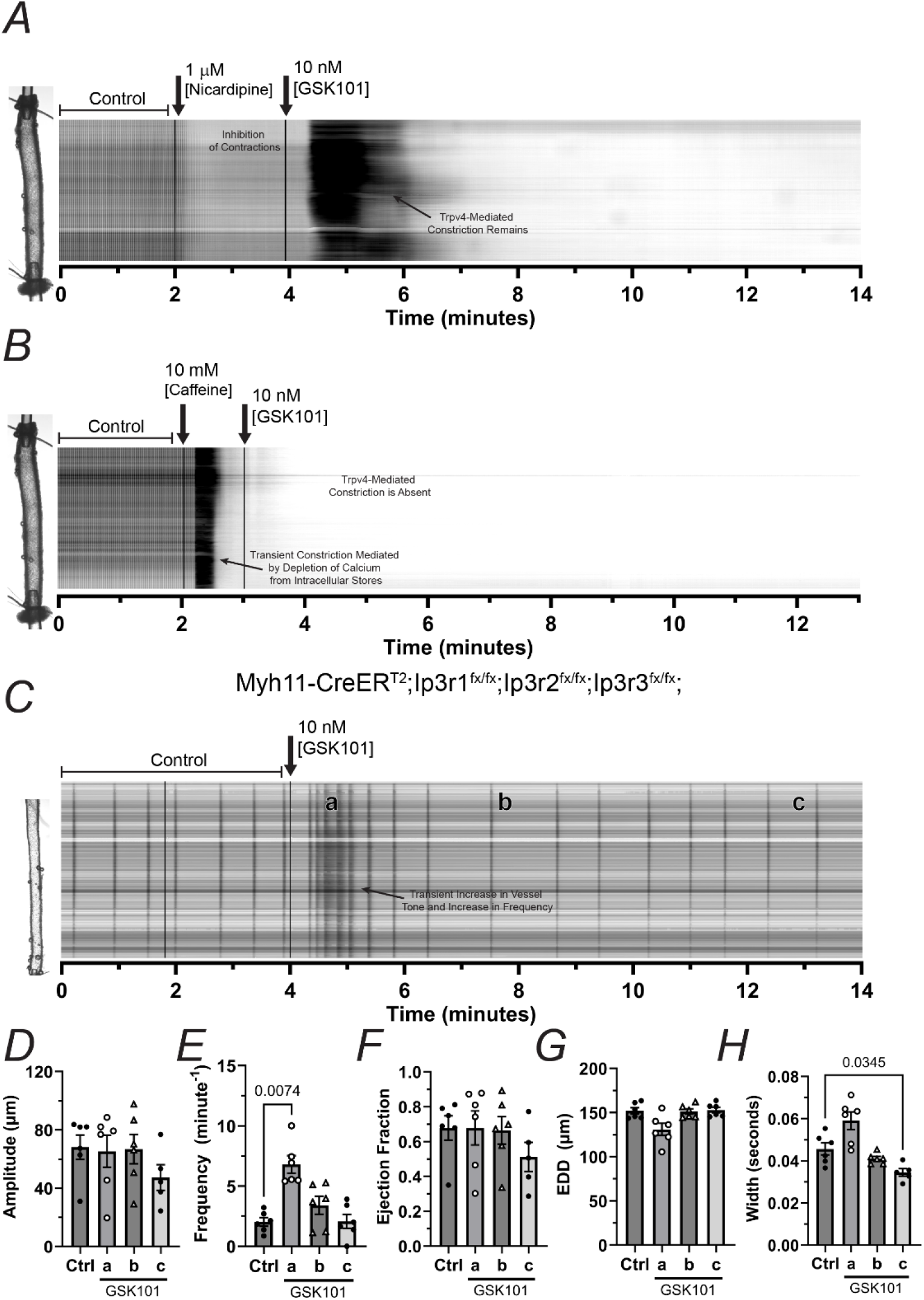
Trpv4-initiated constriction mediated by thromboxane a2 requires calcium from intracellular stores and Ip3 receptors. **(A)** Representative STM example depicting the contractile activity of an inguinal axillary vessel from the mouse under control conditions and following stimulation with GSK101 (10 nM) in the presence of the L-type calcium channel antagonist nicardipine (1 µM). Nicardipine inhibited all contractile activity but did not prevent the strong and sustained constriction induced by GSK101; **(B)** Representative STM example depicting the contractile activity of an inguinal axillary vessel from the mouse under control conditions and upon stimulation with GSK101 (10 nM) following depletion of intracellular calcium using caffeine (10 mM). Caffeine induced a strong, transient constriction of the lymphatic vessel and prevented a subsequent constriction when stimulated with GSK101; **(C)** Representative STM example depicting the contractile activity of an inguinal axillary vessel from an Myh11-CreER^T2^;Ip3r1^fx/fx^;Ip3r2^fx/fx^;Ip3r3^fx/fx^ mouse under control conditions and following stimulation with GSK101 (10 nM); (D-H) Summary of lymphatic contractile parameters **(D)** contraction amplitude, **(E)** contraction frequency, **(F)** ejection fraction, **(G)** end diastolic diameter (EDD), and **(H)** contraction width. Ip3-receptor-deficiency prevented the strong constriction and significant impairment of the contractile function induced by GSK101. STMs shown in A-C are representative examples from a total of n=10, 5, and 5 experiments respectively, including both male and female. Values are reported as mean±SEM. Statistical significance was determined using one-way ANOVA corrected for multiple comparisons using a Dunnett test. Significant differences are shown for p values <0.05.

**Figure 13.**
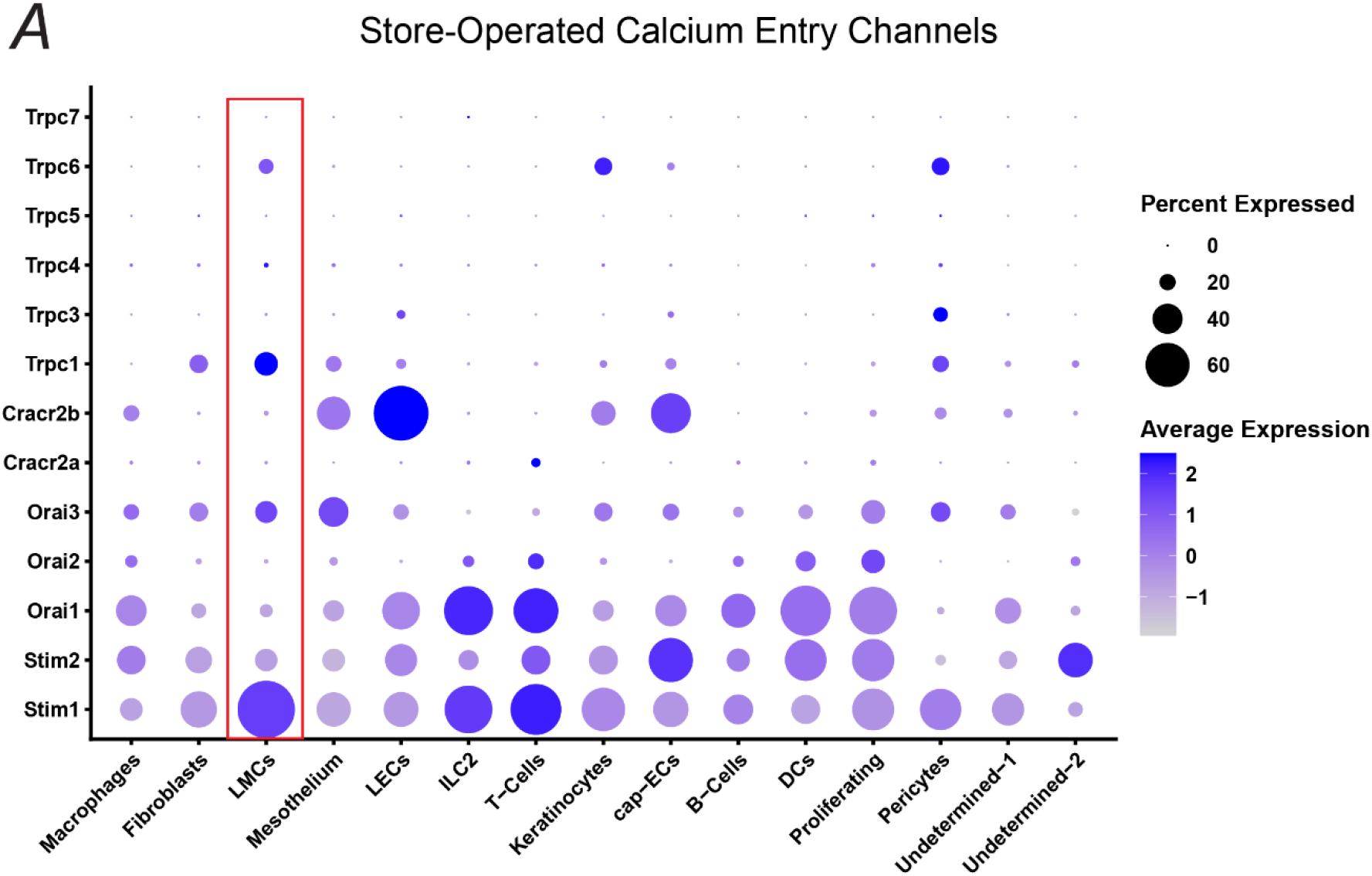
Gene expression for Store Operated Calcium Entry (SOCE)-related channels in collecting lymphatics. (A) DotPlot demonstrating the expression of genes that encode channels known to play a role in SOCE, including Stim1-2, Orai1-3, Cracr2a-b, and Trpc1,3-7. Expression for genes in LMCs is highlighted by a red box. Dominant expressions were observed for *Stim1*, *Orai3*, *Trpc1*, and *Trpc6*. In this DotPlot, the percent of cells expressing a gene of interest within a given class is encoded in the dot size, while the color encodes the average expression level (expressed as *avg_log2FC*) across all cells within a class.

## Discussion

At present, there are no established pharmacological treatments or cures for lymphedema. In developed countries, the largest group of patients afflicted with this disease are cancer-related lymphedema patients who develop the disease following surgical and radiotherapeutic intervention. Clinical evidence has linked tissue fibrosis and inflammation to lymphedema. In a clinical study that included women with stage 2 or 3 unilateral upper-extremity breast cancer-related lymphedema, matched lymphedematous and control biopsies were obtained from each patient. The control biopsy was collected from the unaffected, healthy arm. Biopsies from the extremity affected with lymphedema displayed significantly increased (>3-fold) macrophage infiltration. These observations were recapitulated in a mouse tail model of lymphedema, and the authors further demonstrated that these infiltrating macrophages displayed markers consistent with M2 differentiation^43^.

A major goal of the present study was to determine the regulatory effects of TRPV4 channel activation on the contractile function of isolated, cannulated, and pressurized lymphatic vessels. In addition to lymphatic endothelial cells (LECs), we have identified a subset of macrophages displaying high gene expression of *Trpv4* (∼8-fold higher expression than LECs). Expression of *Trpv4* in LMCs was negligible. Our *controlled* and *spatially defined* scRNAseq data further revealed that these macrophages displayed a transcriptomic profile consistent with that of tissue-resident macrophages (TRMs)^39–42^, differentially expressing *Cd163*, *Folr2*, *Mrc1*, *Ccl8*, *Apoe*, *Cd209f*, *Cd209d*, *Cd209g,* and *Lyve1*. At least half of the macrophages in this subpopulation also expressed another known marker of TRMs, *Timd4*. These observations were confirmed and validated using immunofluorescence and confocal microscopy on isolated lymphatic vessels, where high expression of TRPV4 colocalized with LYVE1-expressing perivascular macrophages.

TRMs are activated by cytokines, including IL-4, IL-10, IL-13, and IL-21, and are known to have anti-inflammatory and angiogenic/lymphangiogenic functions and are involved in tissue remodeling and repair, among others. We demonstrate that signaling through TRPV4 channels leads to the activation of these macrophages and subsequent release of TXA2, leading to major impairment of lymphatic contractility.

To elucidate the mechanism of TRPV4-mediated lymphatic contractile impairment, we utilized pressure myography on isolated lymphatic vessels and automated tracking of changes in inside and outside diameter over time. We generated STMs, which are excellent analyses tools that allow for the representation of an entire video from one experiment (changes in diameter and frequency over time and along an entire lymphatic segment) in a two-dimensional image. Our analyses revealed a multi-phasic effect following the activation of TRPV4 channels. We had anticipated that activation of TRPV4 channels in LECs would result in vasodilation and partial or complete inhibition of spontaneous contractions mediated by nitric oxide. Indeed, this was identified as a secondary component in the observed multi-phasic response which could be prevented by the NOS inhibitor L-NAME; however, our experiments revealed an initial strong and sustained vasoconstrictive component which was prevented in the presence of the TRPV4 selective and potent inhibitor GSK219 and the COX-1,2 inhibitor indomethacin, indicating the involvement of a vasoconstriction-inducing prostanoid(s). Since macrophages and LECs were the two major cell types expressing *Trvp4*, we utilized our scRNAseq dataset to identify these prostanoids by determining the expression of the different prostanoid synthases. In LECs, *Ptgs1* and *Ptges3* displayed the highest expression, while in macrophages, the highest expressed synthases were *Tbxas1* and *Ptges3* (Figure 8C). *Tbxas1* encodes the protein known to catalyze the conversion of prostaglandin H2 to the potent vasoconstrictor TXA2. Given that constriction indicates activation of LMCs, we also looked at expression of the different prostanoid receptors expressed in this cell type. In LMCs, *TXA2r* encoding the G protein-coupled receptor (GPCR) TXA2R was a strong dominant, further suggesting that the strong and sustained constriction observed upon activation of TRPV4 channels is mediated by TXA2. These results were confirmed and functionally validated in isolated collecting lymphatic vessels, where the TRPV4-induced constriction was prevented in the presence of the TXA2R inhibitor, Terutroban (S18886). This constriction remained in the presence of the L-type calcium channel inhibitor, Nicardipine, but was not present in lymphatic vessels from mice deficient in smooth muscle Ip3Rs or in lymphatic vessels from WT animals following depletion of calcium from intracellular stores, suggesting a store-operated calcium entry mechanism initiated by the GPCRs, TXA2Rs.

When assessing the effects of TRPV4 channel activation, our results highlighted both anatomical and sex-based differences. Further experimentation is needed to determine the mechanisms underlying these differences. However, our results suggest that the observed functional differences could be explained by the anatomical and/or sex-based heterogeneity intrinsic to these *Trpv4*-expressing macrophage subpopulations. Although activation of TRPV4 channels resulted in contractile impairment in lymphatic vessels from both male and female mice, lymphatics from male mice displayed a significantly stronger response than those from female mice, this was associated with a decreased density of *Trpv4*-expressing macrophages in lymphatics from female mice.

Importantly, similar functional responses following stimulation of TRPV4 channels were observed in arterioles from the different anatomical areas included in this study. Suggesting that the application and impact of our findings can be, at least to some extent, extrapolated to arterioles. Additional dedicated studies focusing on blood vessels are needed.

In the context of cancer-related lymphedema, and perhaps other forms of secondary lymphedema involving tissue damage/injury, the recruitment of macrophages to the affected area and their anti-inflammatory and pro-lymphangiogenic roles, as well as their role in promoting tissue repair are critical in determining the onset and progression of the disease; however, our results point to detrimental effects to the contractile activity, i.e., pumping capacity, of collecting lymphatic vessels mediated by TRPV4 channels in both LYVE1-expressing macrophages and LECs. Importantly, the functional impairment mediated by TRPV4 channels could extend beyond injured lymphatics, impacting healthy lymphatic vessels located within and in close proximity to the lymphedematous tissues. Our findings suggest that pharmacological targeting of TRPV4 channels in macrophages, for which LYVE1 could be exploited as a surface anchor to increase specificity and reduce off-target effects; or pharmacological targeting of TXA2Rs may offer novel therapeutic strategies to improve lymphatic pumping function and lymph transport in lymphedema.

## Abbreviations

Abbreviations:
| Name | Abbreviation |
| --- | --- |
| Transient receptor potential vanilloid type 4 | TRPV4 |
| Single cell RNA sequencing | scRNAseq |
| Lymphatic endothelial cells | LECs |
| Lymphatic muscle cells | LMCs |
| GSK1016790A (TRPV4 agonist) | GSK101 |
| GSK2193874 (TRPV4 antagonist) | GSK219 |
| Endothelin-1 | ET-1 |
| Physiological saline solution | PSS |
| Prostaglandin | PG |
| Terutroban (TXA2 receptor antagonist) | S18886 |
| Thromboxane A2 | TXA2 |
| Lymphatic vessel endothelial hyaluronan receptor 1 | LYVE1 |

## Acknowledgements

The authors are thankful for the extraordinary help of Dr. Jay Kolls and the “Tulane Center for Translational Research in Infection and Inflammation – NextGen Sequencing Core”, Dr. Georgina L. Dobek and the “Tulane Department of Comparative Medicine”, as well as the “Tulane National Primate Research Center” (RRID: SCR_008167) Pathology Core (RRID: SCR_024606).

