## Supplemental Materials for "TRPV4-Expressing Tissue-Resident Macrophages Regulate the Function of Collecting Lymphatic Vessels via Thromboxane A2 Receptors in Lymphatic Muscle Cells"

**Affiliations:** [1]Department of Pharmacology, Tulane University School of Medicine, New Orleans, LA 70112, [2]Center for Translational Research in Infection and Inflammation, Tulane University School of Medicine, New Orleans, LA 70112, [3]Department of Medical Pharmacology and Physiology, University of Missouri School of Medicine, Columbia, MO 65212, [4]Division of Plastic and Reconstructive Surgery, Department of Surgery, Memorial Sloan Kettering Cancer Center, New York, NY, 10604, USA, [5]Tulane National Primate Research Center, Covington, LA 70433, [6]Immunology Center of Georgia, Medical College of Georgia, Department of Physiology, Augusta University, Augusta, GA 30912, USA.

### Supplemental Figures and Tables

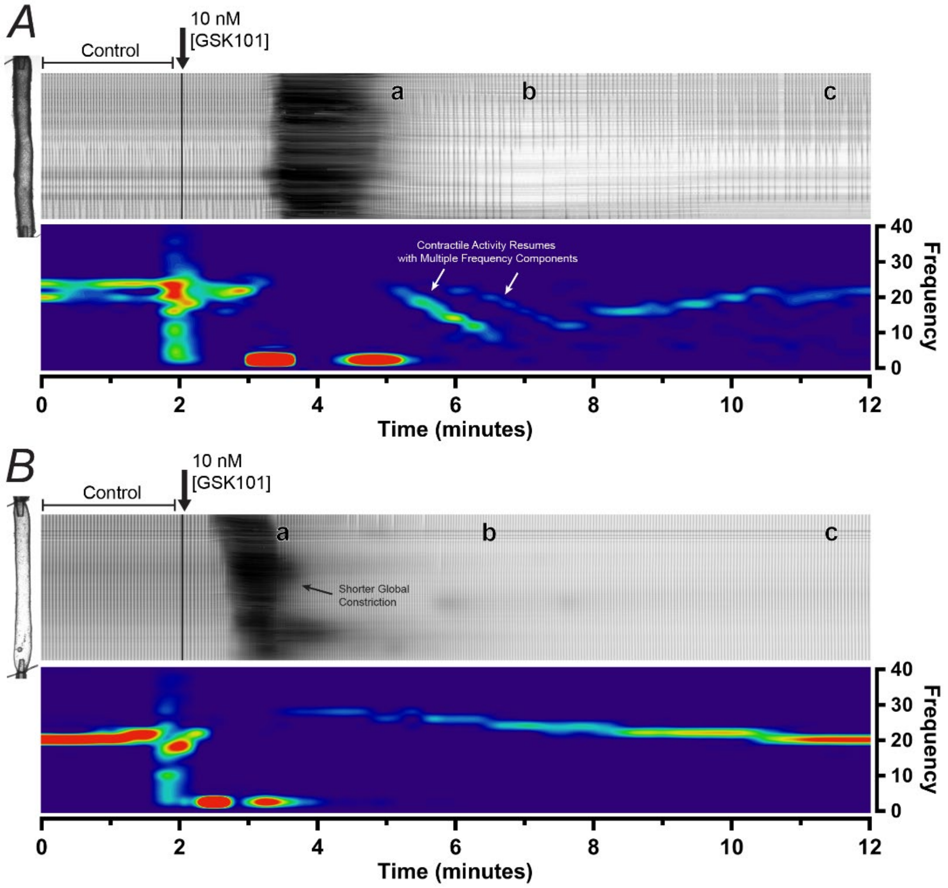


**Supplemental Figure 1. Pharmacological activation of TRPV4 channels results in significant impairment of the contractile function of lymphatic vessels in male and female mice. (A)** Space-time map (STM) and corresponding spectrogram generated from a representative 12-minute video recording of the contractile activity of an inguinal axillary lymphatic vessel (IALV) from male mice. STM displays the contractility of the vessel under control conditions and upon pharmacological activation of TRPV4 channels using GSK101 (10 nM). Sustained, strong, and global constriction was observed in male IALVs following stimulation with GSK101 (dark area between minutes 2-4), then a transient increase in contractile frequency was observed (label “a” on STM), followed by loss of contractile activity, evident by the lack of narrow dark vertical bands (label “b” on STM), and finally contractile activity is partially restored after 10 minutes of drug washout (label “c” on STM). The spectrogram depicts the main frequency components (major frequency components in red over a blue background) as a function of time. **(B)** STMs and corresponding spectrogram generated from a representative example of a 12-minute video recording of the contractile activity of the inguinal axillary lymphatic vessel from a female mouse.

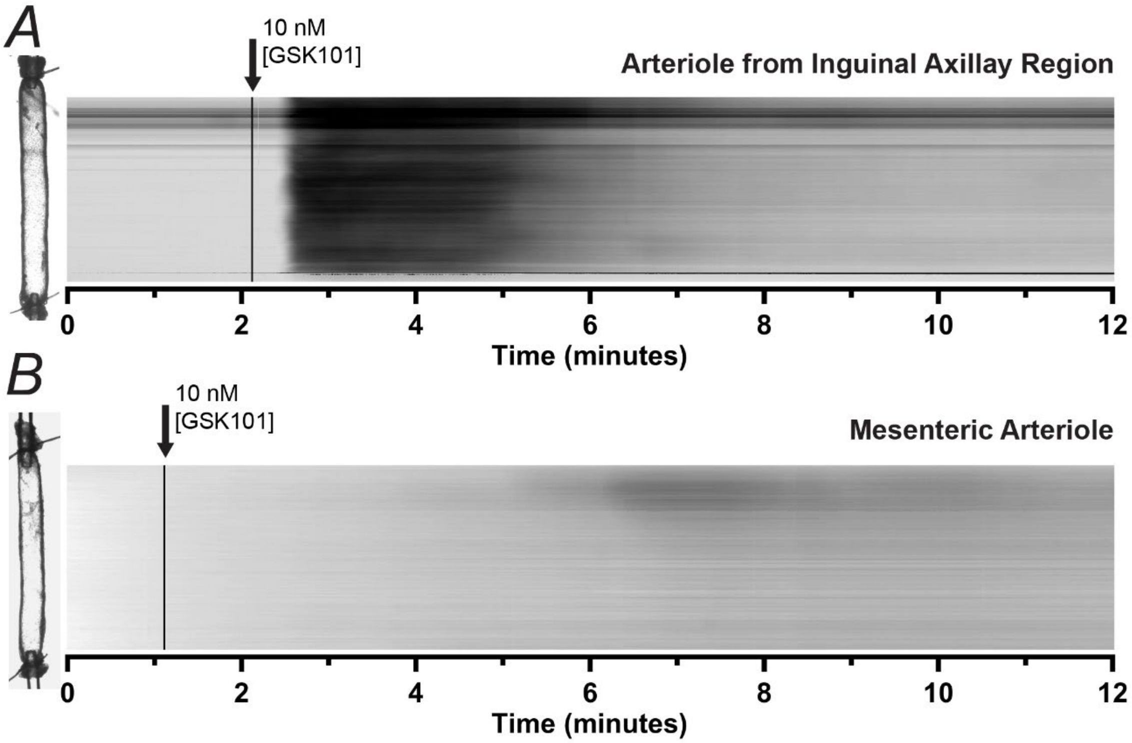


**Supplemental Figure 2. Pharmacological activation of TRPV4 channels results in arterioles from different anatomical regions. (A,B)** STMs generated from representative examples of 12-minute video recordings of the functional response to pharmacological activation of TRPV4 channels using GSK101 in arterioles from the inguinal axillary and mesenteric region. A robust and sustained constriction was observed in arterioles from the inguinal axillary region; however, consistent with mesenteric lymphatics, mesenteric arterioles did not display a robust constriction, only an increase in tone, and local, very low amplitude constriction sites were observed.

**Supplemental Table 1. List of Pharmacological Agents and Antibodies**

| Reagent | Source | Catalog # | Persistent ID / URL |
|---|---|---|---|
| **GSK1016790A** | Tocris | 6433 | https://www.tocris.com/products/gsk-1016790a_6433 |
| **GSK2193874** | Tocris | 5106 | https://www.tocris.com/products/gsk-2193874_5106 |
| **Nicardipine** | Sigma | N7510 | https://www.sigmaaldrich.com/US/en/product/sigma/n7510 |
| **L-Name** | Sigma | N5751 | https://www.sigmaaldrich.com/US/en/product/sigma/n5751 |
| **Indomethacin** | Tocris | 1708 | https://www.tocris.com/products/indomethacin_1708 |
| **S18886** | Tocris | 5568 | https://www.tocris.com/products/s-18886_5568 |
| **U46619** | Tocris | 1932 | https://www.tocris.com/products/u-46619_1932 |
| **Caffeine** | Sigma | C0750 | https://www.sigmaaldrich.com/US/en/product/sial/c0750 |
| **Trpv4 Antibody** | Novus Biologicals | NBP2-41262 | https://www.novusbio.com/products/trpv4-antibody_nbp2-41262 |
| **Lyve1 Antibody** | Novus Biologicals | NBP1-43411 | https://www.novusbio.com/products/lyve-1-antibody-aly7_nbp1-43411 |

| Supplemental Table 2. RT-PCR Primers for Mouse Prostanoid Synthases | | | | |
|---|---|---|---|---|
| **Gene** | **Accession No.** | **Strand** | **Sequence** | **Amplicon Size (bp)** |
| **Ptgds** | NM_008963 | s | GCTCCTTCTGCCCAGTTTTC | 210 |
| | | as | GCCCCAGGAACTTGTCTTGT | |
| **Ptgis** | NM_008968 | s | GTGAGGAGAAGGCCAGGATG | 210 |
| | | as | GGTCTTTATCCCCCGCTGAC | |
| **Ptges** | NM_022415.3 | s | TCGGGGTAAGAGGGCCATTA | 399 |
| | | as | CCAGATTTGCAGCCAGGAGA | |
| **Ptges2** | NM_133783 | s | TGACAGCCGTGGGTAAAGAC | 302 |
| | | as | CAGAAGCAGAGAACTGGGGG | |
| **Ptges3** | NM_019766 | s | GAGAATCCGGCCAGTCATGG | 312 |
| | | as | ATCCAGGCGATGACAACAGC | |
| **Tbxas1** | NM_011539 | s | TTGGGGGACTCACTTGATGG | 311 |
| | | as | ACCATTTCAGGAGGGCCAAG | |
| **Pxrl2b** | NM_025582 | s | GCGACAAGGTACTGTTGCAC | 371 |
| | | as | GTTACAGATGCCTCCGGGTC | |

| Supplemental Table 3. Cell count summary in each sample | | | | | | | | |
|---|---|---|---|---|---|---|---|---|
| **Cell Type** | **Female Mice** | | | | **Male Mice** | | | |
| | **F1** | **F2** | **F3** | **F4** | **M1** | **M2** | **M3** | **M4** |
| **Macrophages** | 665 | 627 | 677 | 671 | 1098 | 980 | 1136 | 1127 |
| **Fibroblasts** | 1461 | 1189 | 836 | 678 | 1453 | 1174 | 1124 | 1033 |
| **LMCs** | 160 | 155 | 194 | 100 | 148 | 233 | 196 | 240 |
| **Mesothelium** | 124 | 95 | 162 | 179 | 22 | 150 | 144 | 105 |
| **LECs** | 42 | 47 | 69 | 69 | 46 | 100 | 84 | 84 |
| **ILC2** | 77 | 67 | 59 | 72 | 27 | 37 | 45 | 59 |
| **T-Cells** | 67 | 104 | 42 | 70 | 27 | 21 | 48 | 49 |
| **Keratinocytes** | 70 | 47 | 72 | 68 | 14 | 22 | 17 | 13 |
| **cap-ECs** | 14 | 33 | 37 | 22 | 30 | 60 | 67 | 44 |
| **B-Cells** | 24 | 68 | 40 | 78 | 22 | 13 | 27 | 28 |
| **DCs** | 41 | 24 | 41 | 27 | 32 | 15 | 29 | 49 |
| **Proliferating** | 24 | 30 | 44 | 22 | 33 | 29 | 11 | 26 |
| **Pericytes** | 19 | 23 | 36 | 26 | 24 | 30 | 25 | 20 |
| **Undetermined-1** | 15 | 13 | 28 | 16 | 10 | 24 | 7 | 10 |
| **Undetermined-2** | 5 | 8 | 12 | 3 | 0 | 0 | 0 | 0 |

| Supplemental Table 4. Quality Control of Single Cell Suspensions for Samples Submitted for scRNAseq | | | | | | | | |
|---|---|---|---|---|---|---|---|---|
| **Parameter** | **Female Mice** | | | | **Male Mice** | | | |
| | **F1** | **F2** | **F3** | **F4** | **M1** | **M2** | **M3** | **M4** |
| **Cell Count** | 262,000 | 156,000 | 199,400 | 240,000 | 166,800 | 134,600 | 264,000 | 156,200 |
| **Cell Viability (%)** | 99.0% | 100.0% | 97.0% | 99.1% | 99.2% | 100.0% | 99.5% | 99.6% |
